# A microRNA expression signature in infant t(4;11) MLL-AF4+ BCP-ALL uncovers novel therapeutic targets

**DOI:** 10.1101/2023.01.12.523775

**Authors:** Camille Malouf, Alasdair Duguid, Giuseppina Camiolo, Leslie Nitsche, Hélène Jakobczyk, Rishi S. Kotecha, Richard A. Anderson, Neil A. Barrett, Owen P. Smith, Katrin Ottersbach

**Affiliations:** Centre for Regenerative Medicine, Institute for Regeneration and Repair, University of Edinburgh, UK, EH16 4UU; Leukaemia Translational Research Laboratory, Telethon Kids Cancer Centre, Telethon Kids Institute, University of Western Australia, Perth, WA, Australia; Curtin Medical School, Curtin University, Perth, WA, Australia; Department of Clinical Haematology, Oncology, Blood and Marrow, Transplantation, Perth Children’s Hospital, Perth, WA, Australia; MRC Centre for Reproductive Health, The Queen’s Medical Research Institute, University of Edinburgh, Edinburgh, UK. EH16 4TJ; Children’s Health Ireland, Department of Haematology, Dublin, Ireland; National Children’s Cancer Service, Children’s Health Ireland, Dublin, Ireland; Trinity College Dublin, Dublin, Ireland

**Author notes:** <u>Contact information for corresponding authors</u>, Dr. Camille Malouf, Professor Katrin Ottersbach, Institute for Regeneration and Repair/Centre for Regenerative Medicine, University of Edinburgh, Edinburgh Bioquarter, 5 Little France Drive, Edinburgh, EH164UU, UK.

**Keywords:** MLL-AF4, leukemia, miR-194, miR-99b, miR-125a-5p, microRNA, drug repurposing, drugs, acetazolamide, tacrolimus, LB-100

## Abstract

Infants and children with MLL-AF4+ leukemia have an urgent need for more efficient and less aggressive therapy. In this study, we studied three microRNAs that are downregulated in MLL-AF4+ B-cell precursor acute lymphoblastic leukemia (BCP-ALL): miR-194, miR-99b and miR-125a-5p. When overexpressed, all three microRNAs impaired the survival of MLL-AF4+ leukemic blasts and the maintenance of MLL-AF4+ BCP-ALL. We identified microRNA target genes responsible for this phenotype that are upregulated in MLL-AF4+ BCP-ALL: CA5B, PPP3CA and PPP2R5C. Using CRISPR-Cas9 and specific inhibitors, we confirmed that CA5B, PPP3CA and PPP2R5C downregulation/inhibition severely compromised the proliferation and survival of MLL-AF4+ leukemic blasts. Importantly, CA5B, PPP3CA and PP2A inhibition by acetazolamide, tacrolimus and LB-100, respectively, showed high toxicity towards MLL-AF4+ leukemic blasts and reduced leukemia burden *in vivo*. This study highlights how the unique microRNA expression signature of patients with MLL-AF4+ BCP-ALL can be used to uncover novel therapeutic avenues and accelerate drug repurposing.

**Statement of significance:** There is an urgent need to identify novel therapeutic avenues for patients with MLL-AF4+ BCP-ALL that are more effective and less aggressive. This study identified three clinically available drugs (acetazolamide, tacrolimus and LB-100) with high and selective toxicity towards MLL-AF4+ leukemic cells.

## Introduction

T(4;11) MLL-AF4 B-cell precursor acute lymphoblastic leukemia (BCP-ALL) is one of the most aggressive malignancies occurring in infants less than one year of age^1^. It is characterized by a poor prognosis and a high frequency of relapse^2–4^. MLL-AF4+ pro-B leukemic blasts display uncontrolled proliferation and harbour an immature pro-B lymphoid phenotype (CD19+CD10-) that retains stem-cell (CD34) and myeloid features (CD33)^5^. Patients with MLL-AF4+ BCP-ALL are also more prone to lineage switching during the course of their therapy and at relapse, suggesting that the cell-of-origin has a dual myeloid/lymphoid potential^6^.

Retrospective analysis of neonatal blood spots and studies in monozygotic twins have confirmed the prenatal origin of infant t(4;11) MLL-AF4+ BCP-ALL^7,8^. The MLL-AF4 fusion oncogene induces a unique epigenetic signature in leukemic blasts that leads to a distinct histone hypermethylation pattern and a strong upregulation of MLL target genes^9–12^. More recent studies have also highlighted the crucial role of the fetal signature and microRNAs in the initiation, maintenance and lineage plasticity of MLL-AF4-driven leukemogenesis^13–18^.

MicroRNAs are aberrantly expressed in leukemia and are crucial to the maintenance of the hematopoietic stem cell pool and blood differentiation^19–23^. In our previous study, we found a low/absent expression of three microRNAs in the leukemic blasts of patients with MLL-AF4+ BCP-ALL: miR-194, miR-99b and miR-125a-5p^14^. These microRNAs have been studied in hematopoiesis and other subtypes of leukemia^21,24^, but their specific role in MLL-AF4+ BCP-ALL remains to be investigated.

In this study, we show that miR-194, miR-99b and miR-125a-5p overexpression impaired the proliferation and survival of leukemic cells *in vitro*, which led to an increased survival of mice with MLL-AF4+ BCP-ALL. Using published gene expression datasets, we identified three target genes upregulated in patients with MLL-AF4+ BCP-ALL that recapitulate the microRNA-mediated phenotype: CA5B (miR-194 target), PPP3CA (miR-99b target) and PPP2R5C (miR-125a-5p target). Knock-out of these genes using CRISPR-Cas9 in leukemic cells, recapitulated the phenotype observed upon their respective microRNA overexpression in all three cases: decreased proliferation and increased apoptosis. Most importantly, there are clinically approved drugs that can inhibit the enzymatic activity of all three genes: acetazolamide (CA5B), tacrolimus (PPP3CA) and LB-100 (PP2A complex). These inhibitors were detrimental to the proliferation and survival of MLL-AF4+ leukemia cell lines and primary cells derived from an infant with MLL-AF4+ BCP-ALL, and all three drugs decreased the MLL-AF4+ leukemia burden *in vivo*. Overall, this study identified three novel tumor suppressor microRNAs in MLL-AF4-driven leukemogenesis (miR-194, miR-99b and miR-125a-5p) that can negatively affect the expression of three oncogenes (CA5B, PPP3CA and PPP2R5C). This led to the identification of novel therapeutic avenues that could be used in the clinic to improve current therapeutic regimens. Hence, the microRNA expression signature of MLL-AF4+ BCP-ALL patients can be used to accelerate the drug discovery process for infant MLL-AF4+ BCP-ALL.

## Results

### MiR-194, miR-99b and miR-125a-5p are downregulated in MLL-AF4+ lymphoid leukemic blasts

To identify candidate tumor suppressor genes in MLL-AF4-driven leukemogenesis, we previously performed microRNA expression profiling in infants and children with t(4;11) MLL-AF4+ BCP-ALL, which led to the identification of 85 differentially expressed microRNAs, including 19 downregulated microRNAs (Figure 1A)^14^. From this list, we selected three microRNA families for further analysis that showed a similar downregulation signature: miR-194, miR-99 family (miR-99a and miR-99b) and miR-125-5p family (miR-125a-5p and miR-125b) (Figure 1B). These microRNAs were among the top 10 downregulated microRNAs^14^. They have been linked to hematopoiesis and/or leukemia^21,24,25^, but have not been studied in the context of MLL-AF4+ BCP-ALL. We specifically selected miR-99b and miR-125a-5p because they are part of the same microRNA cluster. MiR-194, miR-99b and miR-125a-5p are all expressed in human and mouse adult bone marrow (BM) and fetal liver (FL) hematopoietic stem cells (HSC) (Figure 1C-E, Supplemental Figure S1A-C). MiR-194 was also highly expressed in other human FL hematopoietic cells (MPP, LMPP, pre-pro-B, pro-B, B/NK/T lymphoid cells, myeloid cells) compared to MA4+ leukemic blasts (Figure 1C). While, miR-99b and miR-125a-5p were also significantly downregulated in MA4+ leukemic blasts compared to human FL MPP, LMPP and a subset of mature hematopoietic cells, their expression in human FL pre-pro-B and pro-B cells was low/absent (Figure 1D,E).

**Figure 1.**
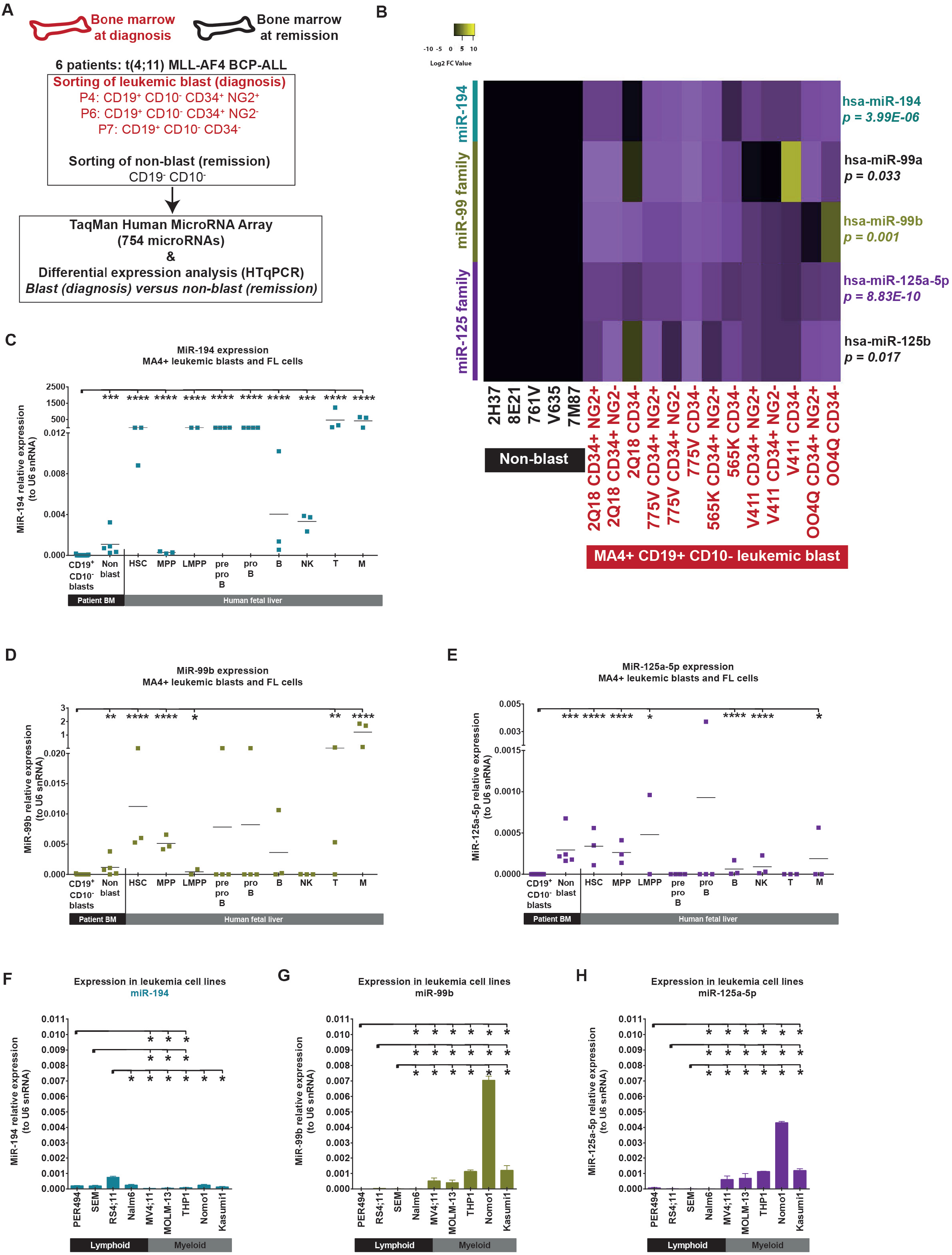
MiR-194, miR-99b and miR-125a-5p are all downregulated in t(4;11) MLL-AF4+ BCP-ALL patients. (A) Experimental design to identify differentially expressed microRNAs in leukemia blasts derived from bone marrow of patients with t(4;11) MLL-AF4+ BCP-ALL. (B) Differential expression analysis of miR-194, miR-99a, miR-99b, miR-125a-5p and miR-125b in CD19+CD10- blasts at diagnosis (NG2+CD34+, NG2-CD34+ and CD34-) compared to non-blasts at remission (CD19-CD10-). Cells were obtained from bone marrow aspirates and data were compared using a limma test. Relative expression of (C) miR-194, (D) miR-99b or (E) miR-125a-5p in MLL-AF4+ leukemic blasts and non-blast MLL-AF4+ BCP-ALL patients alongside human FL HSC, MPP, LMPP, pre-pro-B/pro-B/B/NK/T lymphoid cells and M/myeloid cells. Expression of (F) miR-194, (G) miR-99b and (H) miR-125a-5p in leukemia cell lines. Data is presented as Mean ± SEM and compared using a Mann-Whitney U test with bilateral p-value: p < 0.05 (*), p < 0.01 (**), p < 0.001 (***) and p < 0.0001 (****).

Next, we assessed the expression of miR-194, miR-99b and miR-125a-5p in an array of leukemia cell lines. Mir-194 expression was low in all leukemia cell lines (Figure 1F). Interestingly, miR-99b and miR-125a-5p expression was higher in mixed (MV4;11) and myeloid leukemia cell lines compared to lymphoid cell lines (Figure 1G-H). This correlates with previous studies highlighting their role in myeloid leukemia^25–27^. In agreement with primary patient results, all three microRNAs also had a low/absent expression in SEM and RS4;11 MLL-AF4+ BCP leukemia cell lines compared to known oncogenic co-drivers (miR-128a and miR-130b) (Supplemental Figure S1D,E)^14^. Together, these results highlight the downregulation of miR-194, miR-99b and miR-125a-5p in MLL-AF4+ leukemic cells with a pro-B phenotype.

### MiR-194, miR-99b and miR-125a-5p overexpression has a negative impact on the fitness of MLL-AF4+ lymphoid leukemic blasts

To gain a better understanding of the role of miR-194, miR-99b and miR-125a-5p in the maintenance of MLL-AF4+ leukemic cells with a pro-B phenotype, we first conducted functional validation in SEM cells which are derived from a child with MLL-AF4+ BCP-ALL. This was done by overexpressing each microRNA individually either through the use of a microRNA mimic (transfection) or a lentiviral vector (transduction) (Figure 2A). The overexpression of each microRNA in transfected SEM cells was confirmed by RT-qPCR (Figure 2B). SEM cells transfected with a microRNA mimic of miR-194, miR-99 or miR-125 all showed reduced proliferation (Figure 2C). This phenotype was recapitulated when overexpressing each microRNA by lentiviral transduction (Figure 2D,E). MiR-194, miR-99b or miR-125a-5p overexpression also led to increased apoptosis in SEM cells (Figure 2F). Importantly, the overexpression of miR-194, miR-99b or miR-125a-5p in NSG-SEM xenografts prolonged the latency of MLL-AF4+ BCP-ALL (Figure 2G,H). These results highlight a novel tumor suppressor role for miR-194, miR-99b and miR-125a-5p in MLL-AF4+ BCP-ALL.

**Figure 2.**
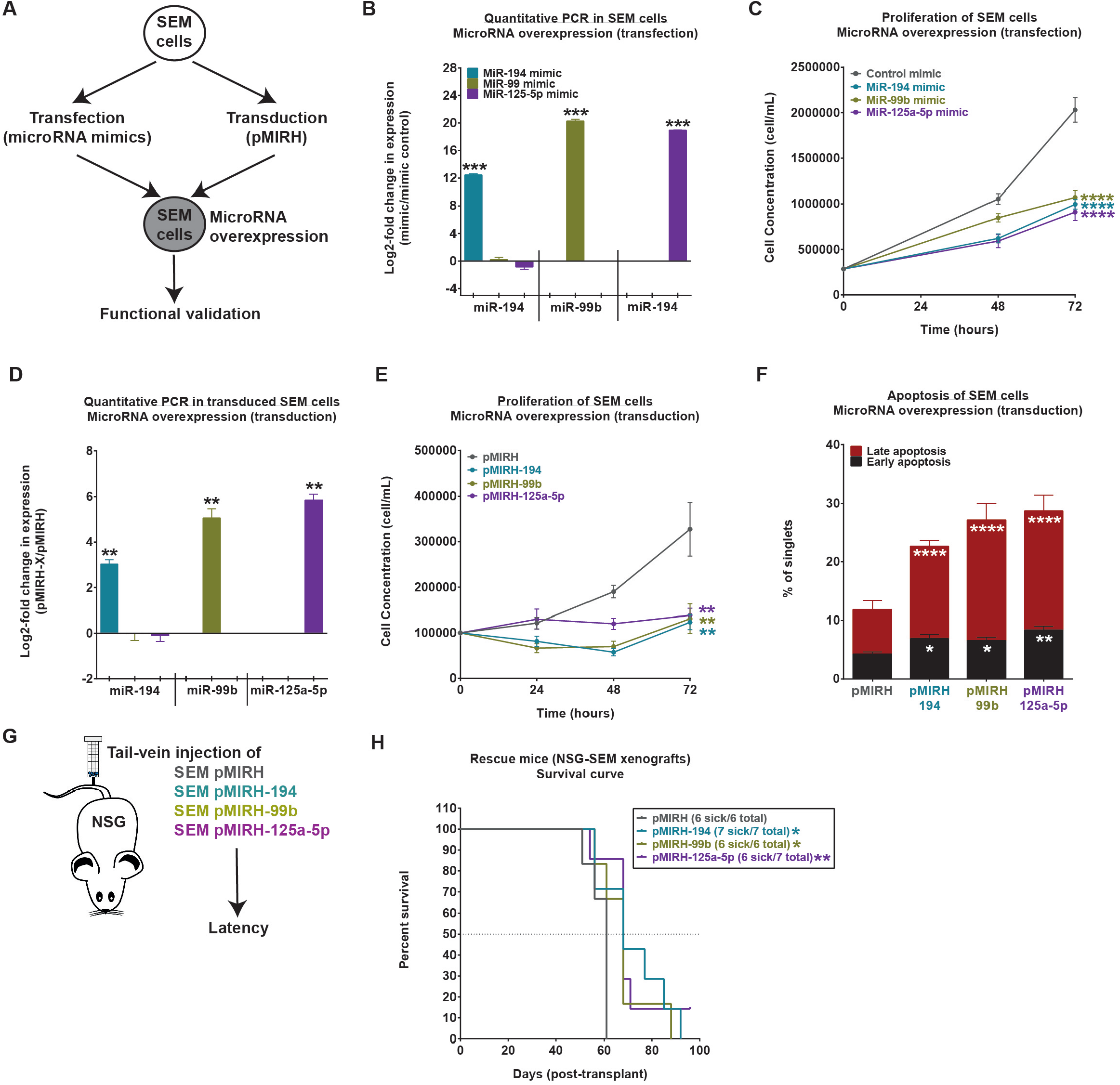
MiR-194, miR-99b and miR-125a-5p overexpression has a negative impact on the fitness of MLL-AF4+ lymphoid leukemic blasts. (A) Experimental layout for functional validation in SEM cells (B) RT-qPCR and (C) proliferation of SEM cells transfected with miRVANA^®^ mimics. (D) RT-qPCR and (E) proliferation of SEM cells transduced with pMIRH lentivirus. (F) Apoptosis of SEM cells overexpressing miR-194, miR-99b or miR-125a-5p through lentiviral transduction. (G) Survival curve of NSG-SEM xenografts that overexpress miR-194, miR-99b or miR-125a-5p. Survival differences were assessed using a Gehan-Breslow-Wilcoxon test. Data is presented as Mean ± SEM and compared using a Mann-Whitney U test with bilateral p-value: p < 0.05 (*), p < 0.01 (**), p < 0.001 (***) and p < 0.0001 (****).

### MiR-194, miR-99b and miR-125a-5p increase the latency of Mll-AF4+ pMIRH-128a pro-B ALL

Next, we assessed the impact of miR-194, miR-99b and miR-125a-5p on the maintenance of Mll-AF4+ pMIRH-128a BCP-ALL, a novel syngeneic mouse model that recapitulates infant t(4;11) MLL-AF4+ BCP-ALL^14^. Similar to patients, all three microRNAs are downregulated in MLL-AF4+ pMIRH-128a BCP-ALL^14^. We harvested BM GFP+ leukemic blasts of Mll-AF4+ pMIRH-128a mice and overexpressed each microRNA individually using lentiviral transduction and transplanted them into Ly5.1 HET recipients (Figure 3A,B). Similar to NSG-SEM mice (Figure 2H), Mll-AF4+ pMIRH-128a BCP-ALL mice that overexpress miR-194, miR-99b or miR-125a-5p showed a longer latency compared to control mice (Figure 3C). All mice eventually developed and succumbed to Mll-AF4+ pro-B ALL with hepatosplenomegaly (Figure 3D,E) and a high GFP chimerism in the bone marrow, spleen, peripheral blood, liver and lungs (Figure 3F, Supplemental Figure S2A-D). Interestingly, all three microRNAs led to a reduction in LSK IL7R+ leukemia propagating cells in the bone marrow (Figure 3G), especially cells that were CKIT^high^ (Figure 3H, Supplemental Figure S2E). Finally, there were more apoptotic cells in Mll-AF4+ pMIRH-128a+ BCP-ALL mice that overexpressed miR-194, miR-99b and miR-125a-5p (Figure 3I). similar to our observation in SEM cells (Figure 2F). We also confirmed the maintained expression of all three microRNAs in Mll-AF4+ pMIRH-128a+ rescue mice (Supplemental Figure S2F). Overall, these results corroborate with our findings in SEM cells: the overexpression of miR-194, miR-99b and miR-125a-5p impairs the maintenance of MLL-AF4+ BCP-ALL.

**Figure 3.**
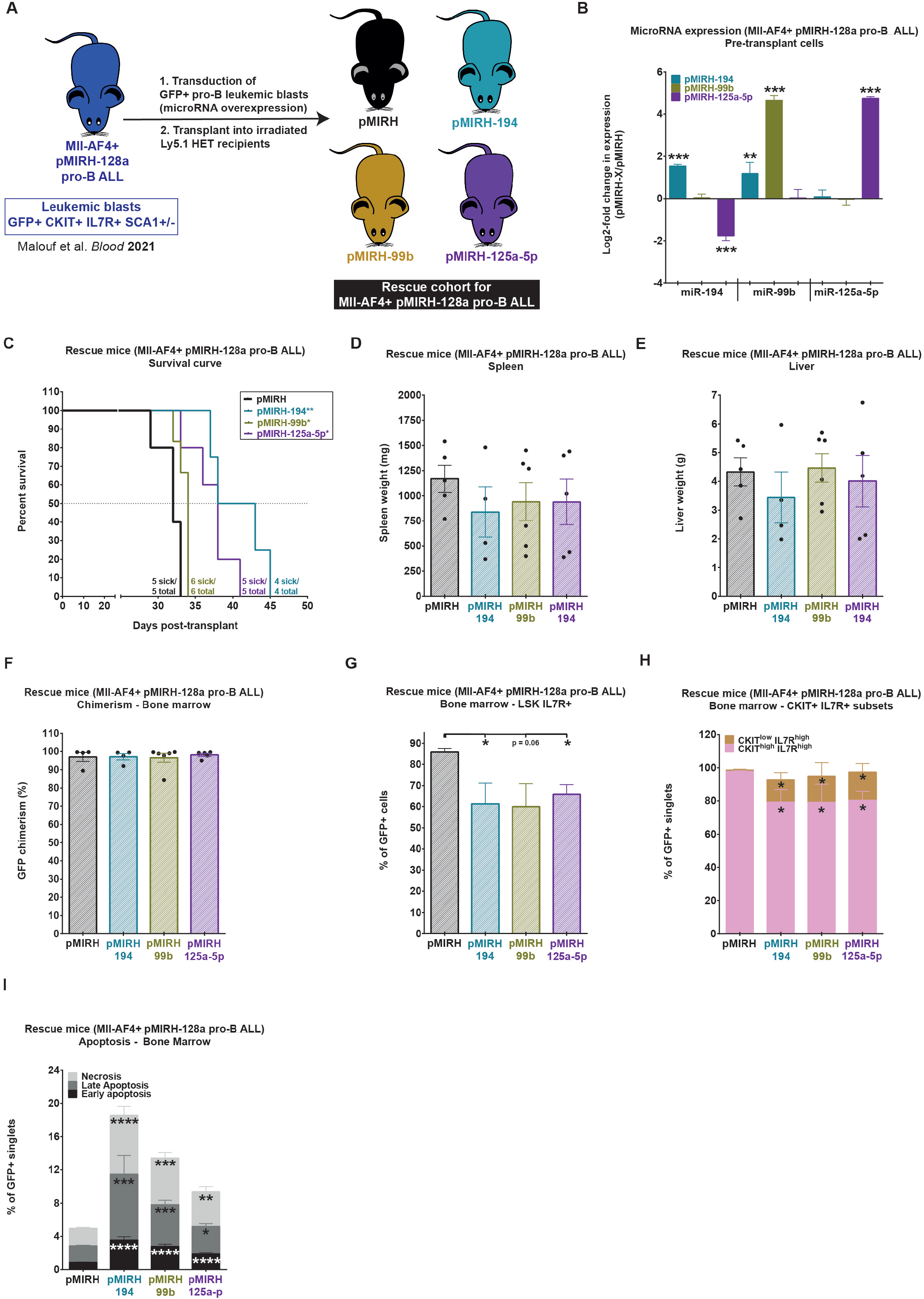
MiR-194, miR-99b and miR-125a-5p increase the latency of Mll-AF4+ pMIRH-128a pro-B ALL. (A) Experimental design. MiR-194, miR-99b and miR-125a-5p are overexpressed in GFP+ Mll-AF4+ pMIRH-128a pro-B leukemic blasts using a lentiviral transduction (pMIRH), followed by transplantation into Ly5.1 HET recipients. (B) RT-qPCR in transduced Mll-AF4+ pre-transplant cells to confirm the overexpression of miR-194, miR-99b or miR-125a-5p. (C) Survival curve of Mll-AF4+ pMIRH-128a mice that overexpress miR-194, miR-99b or miR-125a-5p in GFP+ Mll-AF4+ pMIRH-128a pro-B leukemic blasts. Survival differences were assessed using a Gehan-Breslow-Wilcoxon test. (D) Spleen weight, (E) liver weight and (F) bone marrow GFP chimerism of Mll-AF4+ pMIRH-128a BCP-ALL mice that overexpress miR-194, miR-99b or miR-125a-5p. Proportion of (G) LSK IL7R+ and (H) CKIT^high^IL7R+/CKIT^low^IL7R+ leukemic blasts in the bone marrow of Mll-AF4+ pMIRH-128a BCP-ALL mice that overexpress miR-194, miR-99b or miR-125a-5p. (I) Apoptosis of GFP+ cells from the bone marrow of Mll-AF4+ pMIRH-128a BCP-ALL mice that overexpress miR-194, miR-99b or miR-125a-5p (leukemia). Data is presented as Mean ± SEM and compared using a Mann-Whitney U test with bilateral p-value: p < 0.05 (*), p < 0.01 (**), p < 0.001 (***) and p < 0.0001 (****).

### MiR-194, miR-99b and miR-125a-5p act through their downstream targets CA5B, PPP3CA and PPP2R5C

Next, we focused on identifying the microRNA target genes downstream of miR-194, miR-99b or miR-125a-5p that are upregulated in leukemic blasts derived from patients with MLL-AF4+ BCP-ALL. We cross-referenced the predicted targets of all three microRNAs with patient expression datasets^28,29^ and identified three genes that are upregulated in patients and that have available drugs in the clinic: CA5B (predicted target of miR-194, inhibited by acetazolamide), PPP3CA (predicted target of miR-99b, inhibited by tacrolimus) and PPP2R5C (predicted target of miR-125a-5p, inhibited by LB-100) (Figure 4A). We confirmed that these are upregulated in patients with MLL-AF4+ BCP-ALL (Figure 4B-D). *CA5B* was also upregulated in MLL-AF4+ leukemic blasts compared to FL and cord blood pro-B lymphoid cells (Figure 4B, Supplemental Figure S3A). *PPP3CA* and *PPP2R5C* expression in FL pro-B lymphoid cells was similar to MLL-AF4+ leukemic blasts, but was significantly higher compared to cord blood pro-B lymphoid cells (Figure 4C,D, Supplemental Figure S3B,C). This could be linked to the fetal signature maintained in MLL-AF4+ BCP-ALL patients^17,18^. Finally, we observed high expression of *CA5B, PPP3CA* and *PPP2R5C* in MLL-AF4+ leukemic cell lines with a pro-B phenotype (PER-494, SEM and RS4;11) (Supplemental Figure S3D-F). Importantly, these respective target genes were downregulated in Mll-AF4+ pMIRH-128a leukemic blasts transduced with pMIRH-194, pMIRH-99b and pMIRH-125a-5p (Supplemental Figure S3G-I).

**Figure 4.**
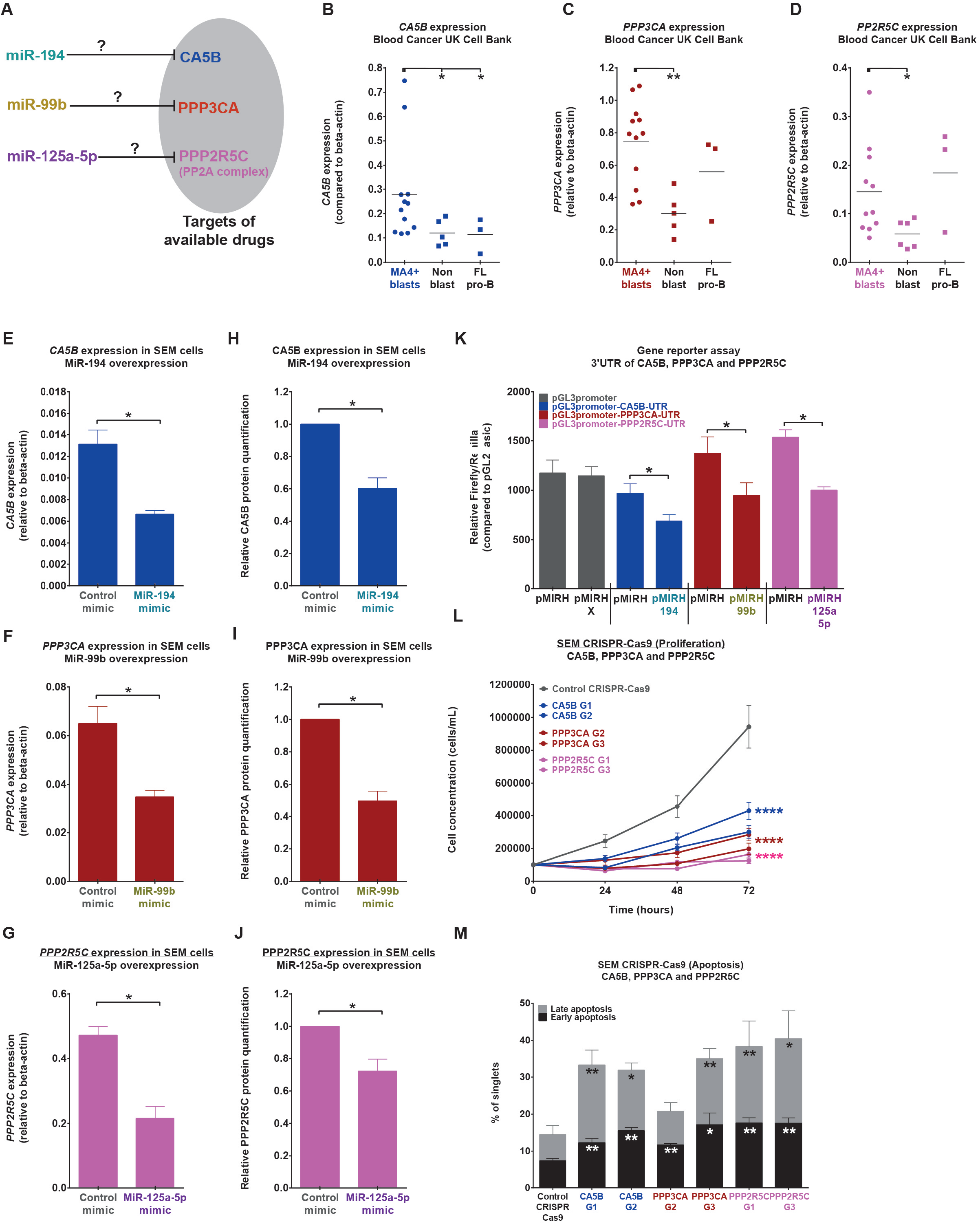
MiR-194, miR-99b and miR-125a-5p act through their downstream targets CA5B, PPP3CA and PPP2R5C. (A) CA5B, PPP3CA and PPP2R5C are predicted targets of miR-194, miR-99b and miR-125a-5p, respectively. (B) *CA5B*, (C) *PPP3CA* and (D) *PPP2R5C* expression in leukemic blasts, non-blasts from MLL-AF4+ BCP-ALL patients and normal FL pro-B lymphoid cells. RT-qPCR of (E) *CA5B*, (F) *PPP3CA* and (G) *PPP2R5C* in SEM cells that overexpress miRVANA^®^ mimics. Western blot of (H) *CA5B*, (I) *PPP3CA* and (J) *PPP2R5C* in SEM cells that overexpress miRVANA^®^ mimics. (K) Luciferase assay in 293T cells using pGL3promoter-UTR vectors of CA5B, PPP3CA and PPP2R5C to assess microRNA-mediated regulation. (L) Proliferation and (M) apoptosis of SEM CRISPR-Cas9 cells knocked-out with CA5B, PPP3CA and PPP2R5C. Data is presented as Mean ± SEM and compared using a Mann-Whitney U test with bilateral p-value: p < 0.05 (*), p < 0.01 (**), p < 0.001 (***) and p < 0.0001 (****).

To confirm the direct regulation of miR-194-CA5B, miR-99b-PPP3CA and miR-125a-5p-PPP2R5C, we conducted gene expression analysis in SEM cells transfected with microRNA mimics (Figure 2B). The overexpression of all three microRNAs led to a significant downregulation of their respective target genes at the RNA (RT-qPCR) and protein level (western blot) (Figure 4E-J, Supplemental Figure S3J). To confirm direct targeting, we cloned the 3’UTR region of each gene that is predicted to be recognized by their respective microRNAs downstream of the Firefly luciferase gene of the pGL3-promoter vector. The significant reduction of the relative luciferase activity upon co-transfection of each UTR with its respective microRNA confirmed the direct regulation between miR-194-CA5B, miR-99b-PPP3CA, and miR-125a-5p-PPP2R5C (Figure 4K). Finally, we assessed the biological consequences of CA5B, PPP3CA or PPP2R5C knock-out in SEM cells using CRISPR-Cas9. SEM cells were co-transduced with a gRNA lentivirus (BFP) and Cas9 lentivirus (GFP). We confirmed the downregulation of CA5B (RNA and protein), PPP3CA (only RNA) and PPP2R5C (RNA and protein) (Supplemental Figure S3K-M). SEM cells that were knocked-out with CA5B, PPP3CA or PPP2R5C all showed a significant decrease in cell proliferation and increased cell death (Figure 4L,M). These results highlight a direct link between three microRNAs that act as tumor suppressors (miR-194, miR-99b and miR-125a-5p) and three oncogenes (CA5B, PPP3CA and PPP2R5C) in MLL-AF4+ BCP-ALL.

### Acetazolamide, tacrolimus and LB-100 impair the survival of MLL-AF4+ pro-B leukemic blasts while having minimal effect on normal FL and BM mouse LSK

Acetazolamide and tacrolimus are commonly used medications: acetazolamide for the treatment of altitude sickness and glaucoma, and tacrolimus as an immunosuppressant to prevent organ transplant rejection^30,31^. LB-100 is currently undergoing clinical trials in small cell lung cancer and recurrent glioblastoma (NCT04560972, NCT03027388). Acetazolamide, tacrolimus and LB-100 inhibit the enzymatic activity of CA5B, PPP3CA and PPP2R5C, respectively (Figure 5A). We sought to determine their effect on MLL-AF4+ leukemic blasts (cell lines and patient cells) and on normal mouse LSK cells derived from 1-month-old bone marrow and E14 FL (Figure 5A). First, we exposed MLL-AF4+ leukemic cell lines (SEM, RS4;11, PER-494 and MV4;11) to all three drugs separately (10 μM), which led to a significant decrease in cell proliferation (Figure 5B, Supplemental Figure S4A-C). All three drugs led to an increase in apoptosis when incubated with SEM cells (Figure 5C), which is reminiscent of the increased apoptotic response upon miR-194, miR-99b or miR-125a-5p overexpression (Figure 2F). Similarly, we treated MLL-AF4+ leukemic cells derived from an infant diagnosed with t(4;11) MLL-AF4+ BCP-ALL and assessed the apoptotic response. Acetazolamide, tacrolimus and LB-100 all led to an increased apoptotic response in primary patient cells (Figure 5D,E).

**Figure 5.**
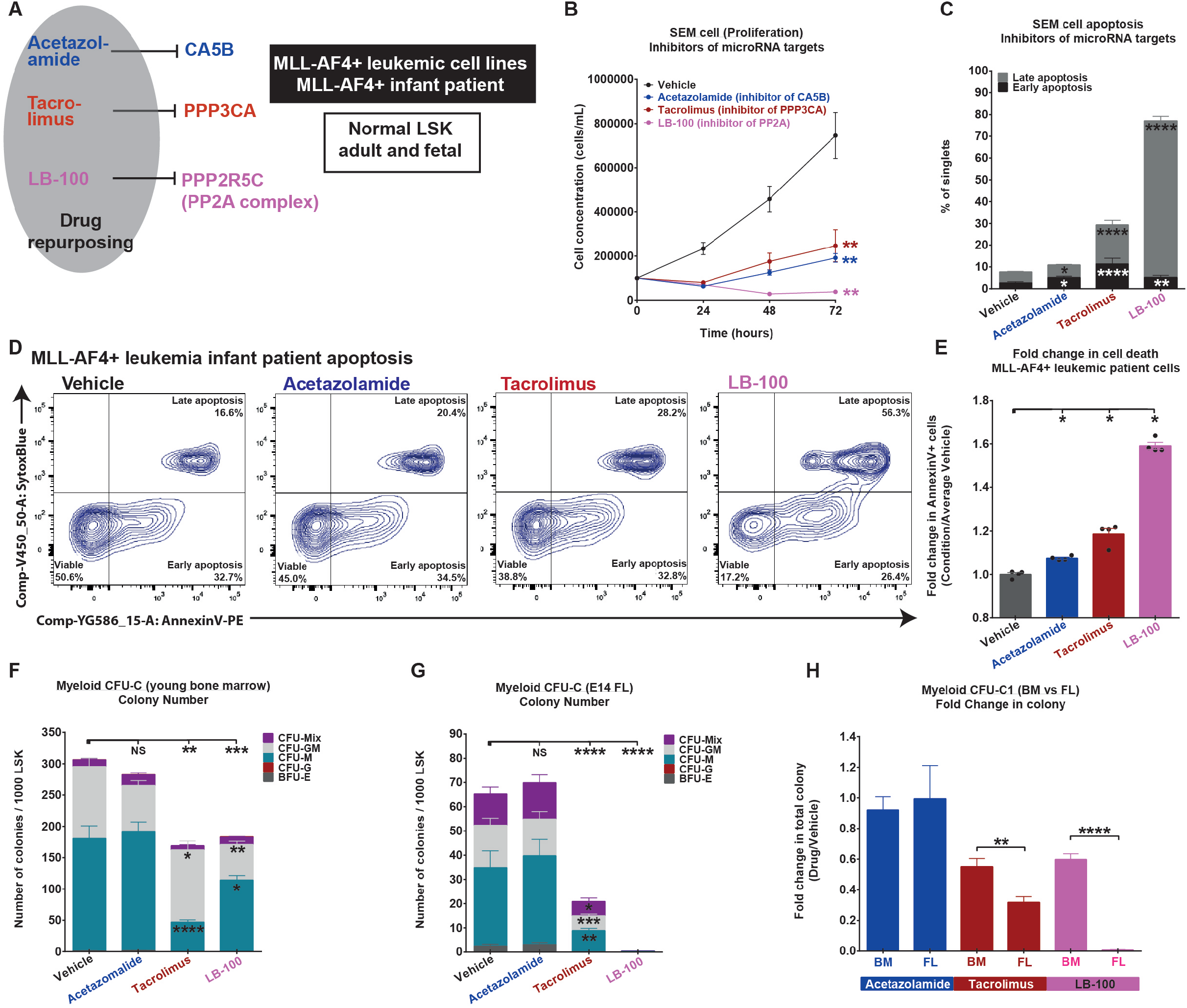
Acetazolamide, tacrolimus and LB-100 impair the survival of MLL-AF4+ pro-B leukemic blasts while having minimal effect on normal FL and BM mouse LSK. (A) Experimental design to assess drug effect on pro-B leukemic cells (SEM and infant MLL-AF4+ patient) and normal young BM and FL mouse LSK. (B) Proliferation and (C) apoptosis of SEM cells exposed to acetazolamide, tacrolimus and LB-100 (10 μM). (D) Apoptosis and (E) fold chance in AnnexinV+ cells of leukemic blasts derived from an MLL-AF4+ BCP-ALL infant patient that were treated with acetazolamide, tacrolimus and LB-100 for 24h (10 μM). Myeloid CFU-C assay of mouse LSK derived from (F) young BM and (G) E14 FL with acetazolamide, tacrolimus and LB-100 (10 μM). (H) Fold change in myeloid CFU-C colony number in mouse young BM and E14 FL LSK exposed to acetazolamide, tacrolimus and LB-100 at 10 μM compared to vehicle. Data is presented as Mean ± SEM and compared using a Mann-Whitney U test with bilateral p-value: p < 0.05 (*), p < 0.01 (**), p < 0.001 (***) and p < 0.0001 (****).

Next, we exposed normal mouse LSK cells derived from young bone marrow (1-month-old) and E14 FL to all three drugs and conducted colony-forming assays to determine their effect on hematopoietic clonogenic potential. Upon the first plating, acetazolamide did not alter the colony-forming capacity of young bone marrow and E14 FL mouse LSK cells (Figure 5F,G). Tacrolimus and LB-100 both led to a significant decrease in hematopoietic colony-forming ability in both young bone marrow and E14 FL mouse LSK cells (Figure 5F,G), but this effect was significantly stronger in E14 FL LSK (Figure 5H). Finally, preliminary results suggest that all three drugs increased apoptosis in human FL CD34+ cells, which have a more proliferative phenotype and an expression signature similar to MLL-AF4+ BCP-ALL patients (Supplemental Figure S4D)^17,18^. Overall, these results show that acetazolamide, tacrolimus and LB-100 can negatively affect the proliferation and survival of MLL-AF4+ leukemic blasts *in vitro*, while they display less toxicity on normal young LSK compared to FL LSK. These features (strong anti-leukemic activity, little/no toxicity towards normal LSK in vitro) make these drugs attractive candidates for the treatment of infants and children with t(4;11) MLL-AF4+ BCP-ALL.

### Acetazolamide, tacrolimus and LB-100 decrease the leukemia burden in MLL-AF4+ BCP-ALL mice

Given that acetazolamide, tacrolimus and LB-100 severely impaired the survival of MLL-AF4+ leukemic blasts *in vitro*, we next evaluated if they could decrease leukemia burden *in vivo*. For the first drug study, we used NSG-SEM BCP-ALL mice to assess each drug separately to determine their effect on MLL-AF4+ BCP-ALL burden and also their overall toxicity. Once human leukemic cells were detected in the peripheral blood (~1%), we initiated a 12-day treatment period (Figure 6A). All NSG-SEM BCP-ALL mice were sacrificed the day after the last drug injection to assess the overall leukemia burden in the peripheral blood, bone marrow, spleen and liver. Firstly, during the course of the treatment, we observed a reduced percentage of human MLL-AF4+ leukemic blasts in the peripheral blood of all NSG-SEM treated mice (Figure 6B). This effect became noticeable at day 8, which is when the leukemia burden of NSG-SEM untreated mice increased exponentially. Notably, 2/4 NSG-SEM vehicle mice developed full-blown leukemia already at day 9 of treatment (Supplemental Figure S5A). This reduced percentage of human MLL-AF4+ leukemic blasts in the peripheral blood of all NSG-SEM treated mice was maintained at day 12 (Figure 6B). NSG-SEM vehicle mice showed a significant decrease in their body weight over the course of treatment (Supplemental Figure S5A). NSG-SEM Acetazolamide mice showed no significant weight loss, while NSG-SEM Tacrolimus and NSG-SEM LB-100 mice showed significant weight loss before the exponential phase of MLL-AF4+ BCP-ALL (Supplemental Figure S5B-D).

**Figure 6.**
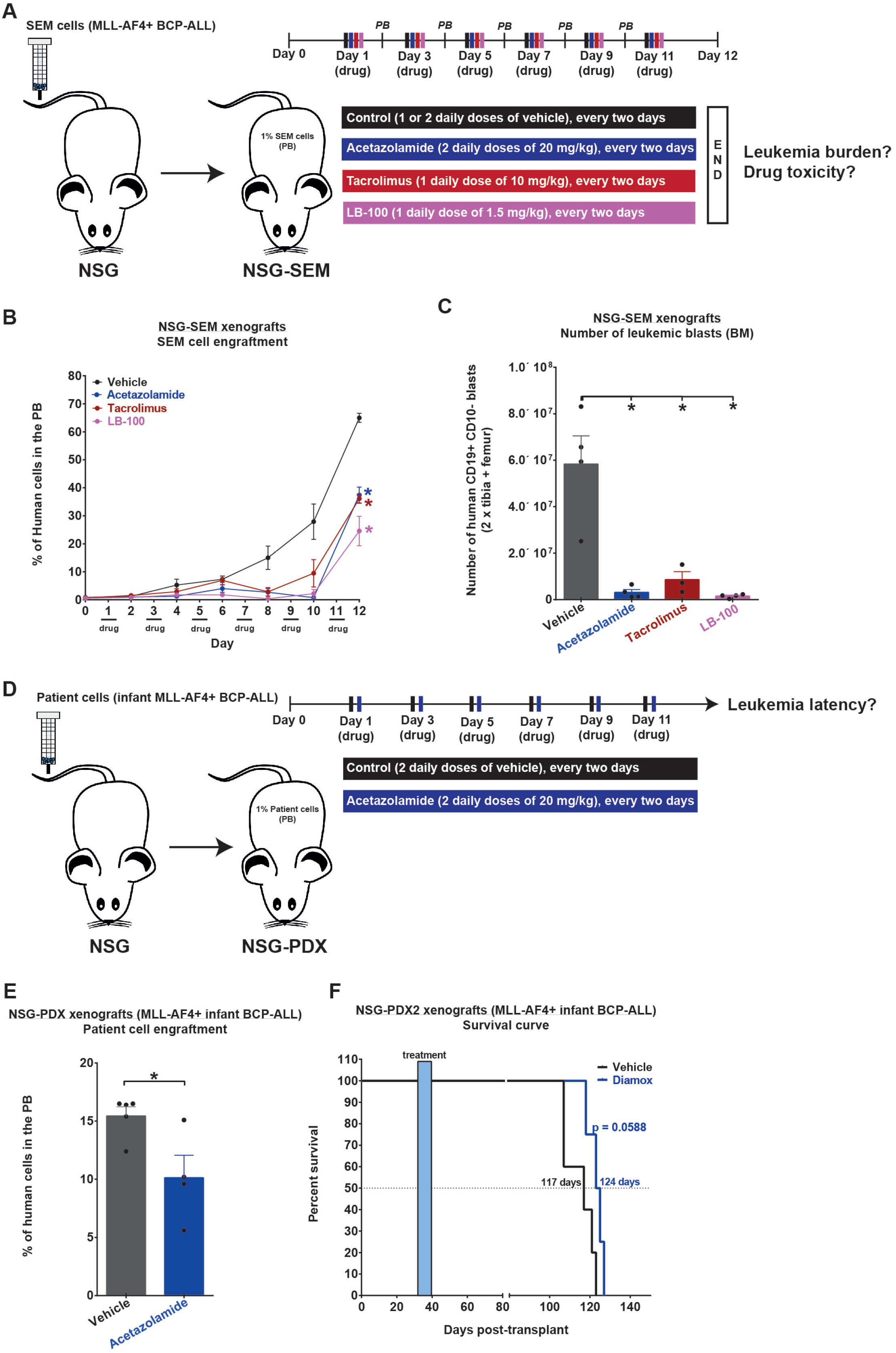
Acetazolamide, tacrolimus and LB-100 impair the maintenance of MLL-AF4+ BCP-ALL. (A) Experimental design of the drug study to assess the effect of acetazolamide, tacrolimus and LB-100 in the maintenance of MLL-AF4+ BCP-ALL in NSG-SEM mice. The treatment was initiated when 1% human cells were detected in the peripheral blood of NSG-SEM mice. (B) Human cell contribution in the peripheral blood over the course of the 12-day treatment. (C) Absolute number of SEM cells in the bone marrow of NSG-SEM mice (vehicle and drug) at day 12. (D) Experimental design of the drug study to assess the effect of acetazolamide on the survival of NSG-PDX MLL-AF4+ BCP-ALL (patient cells, infant MLL-AF4+ BCP-ALL). (E) Human cell contribution in the peripheral blood 5 days after the end of the treatment. (F) Survival curve of NSG-PDX vehicle and NSG-PDX acetazolamide mice. Survival differences were assessed using a Gehan-Breslow-Wilcoxon test. Data is presented as Mean ± SEM and compared using a Mann-Whitney U test with bilateral p-value: p < 0.05 (*), p < 0.01 (**), p < 0.001 (***) and p < 0.0001 (****).

Next, we quantified MLL-AF4+ leukemia burden in the bone marrow, spleen and liver of all mice at day 12 (the day after the last injection). Strikingly, we observed 6-10 times fewer hCD19+hCD10- leukemic cells in the bone marrow of all NSG-SEM treated mice (Figure 6C). This trend was also observed in the spleen of NSG-SEM Acetazolamide mice (Supplemental Figure S5E). The liver size was similar between NSG-SEM vehicle and treated mice (Supplemental Figure S5F). This suggests that the drug-sensitivity of MLL-AF4+ leukemic cells could be variable between tissues.

Given that acetazolamide had the most anti-leukemia activity and the least toxicity amongst all three drugs, we decided to assess its efficacy on the survival of NSG-PDX MLL-AF4+ BCP-ALL mice (Figure 6D). These NSG mice received leukemic cells from an infant MLL-AF4+ BCP-ALL patient (Figure 6D). Acetazolamide reduced the human leukemia burden in the peripheral blood (Figure 6E). NSG-PDX mice treated with acetazolamide also survived for a longer period (p = 0.07) compared to NSG-PDX vehicle mice (124 days *vs* 117 days). Given that the treatment was short and administered two months before the development of full-blown leukemia, these results suggest that acetazolamide has anti-leukemic activity. Overall, this study has identified three drugs that could be beneficial for the treatment of infant MLL-AF4+ BCP-ALL patients.

## Discussion

The aim of this study was to identify novel therapeutic avenues for MLL-AF4+ BCP-ALL patients using their unique microRNA expression signature. Given that microRNAs are negative regulators of gene expression, we selected three microRNAs that are downregulated in infant MLL-AF4+ BCP-ALL patients: miR-194, miR-99b and miR-125a-5p^14^. MiR-194 can act as a tumor suppressor in myeloid leukemic cells^24^, while the miR-99b/let7e/miR-125a-5p cluster is important for maintaining the hematopoietic stem cell pool in human and mice^21,32,33^. MiR-99 and miR-125 families can also provide skewing towards the myeloid lineage and contribute to the development of myeloid malignancies. It is worth mentioning that miR-125a-5p overexpression in FL Mll-AF4+ LSK pre-leukemic cells caused myeloid malignancy in about 50% of recipients, which suggests that it can favor myeloid lineage in MLL-AF4+ hematopoietic stem and progenitor cells. Notably, miR-99 family members are usually downregulated in ALL patients compared to AML patients^34^. Therefore, we decided to investigate if the downregulation of miR-194, miR-99b and miR-125a-5p in infant MLL-AF4+ BCP-ALL was linked to an unknown tumor suppressive role in this specific leukemia subtype.

We conducted extensive functional validation *in vitro* and *in vivo* using both mouse and human models of infant and pediatric MLL-AF4+ BCP-ALL. First, we overexpressed all three microRNAs separately in SEM leukemic cells which are derived from a child with MLL-AF4+ BCP-ALL. Similar to patients, SEM cells have a very low/absent expression of miR-194, miR-99b and miR-125a-5p. Their overexpression decreased proliferation and increased apoptosis in SEM cells. All three microRNAs also prolonged the survival of MLL-AF4+ BCP-ALL mice (NSG-SEM and Mll-AF4+ pMIRH-128a pro-B ALL). These results suggest that microRNA function can be similar between leukemia subtypes (miR-194)^24^, but can also differ (miR-99b and miR-125a-5p)^25,26^.

We next aimed to identify relevant target genes for each microRNA that are upregulated in infants with MLL-AF4+ BCP-ALL, which could be treated by novel drugs that are readily available for clinical use. We focused on CA5B (miR-194 target), PPP3CA (miR-99b target) and PPP2R5C (miR-125a-5p target) that can be inhibited by acetazolamide, tacrolimus and LB-100, respectively. CA5B is a mitochondrial carbonic anhydrase that catalyses the reversible hydration of carbon dioxide, which is essential to many biological functions including pH balance, cell metabolism and enzymatic activity^35^. PPP3CA is a protein phosphatase that responds to calcium signalling to activate specific gene expression programs^36^, while the PP2A phosphatase complex which comprises the PPP2R5C subunit is a major signalling hub in both normal and malignant cells^37^. All three genes are overexpressed in MLL-AF4+ BCP-ALL patients and showed significant downregulation upon their respective microRNA overexpression in SEM cells and pro-B leukemic cells derived from Mll-AF4+ pMIRH-128a mice. The gene reporter assay using the 3’UTR confirmed the direct microRNA-mediated regulation of these target genes. The knock-out of CA5B, PPP3CA or PPP2R5C using CRISPR-Cas9 in SEM cells also recapitulated the phenotype observed upon their respective microRNA overexpression. We also noted a severe decrease in the proliferation and survival of MLL-AF4+ leukemic cells (SEM, PER-494, RS4;11 and MV4;11) and primary cells derived from an infant diagnosed with MLL-AF4+ BCP-ALL upon exposure to acetazolamide, tacrolimus or LB-100 *in vitro*. Notably, acetazolamide showed very little toxicity towards normal young BM and E14 FL LSK cells. Tacrolimus and LB-100 showed a moderate toxicity towards normal LSK cells, but to a less extent than towards Mll-AF4+ leukemic cells. Their effect also appeared stronger in FL LSK cells which have a more proliferative phenotype and a gene expression signature similar to infant MLL-AF4+ BCP-ALL patients^17,18^. Overall, these results show a very good response of MLL-AF4+ leukemic cells to acetazolamide, tacrolimus and LB-100. They also highlight novel pathways that can influence the leukemia microenvironment and cell metabolism (CA5B), but also intracellular signalling and gene expression (PPP3CA and PPP2R5C).

Finally, we assessed the efficiency of acetazolamide, tacrolimus and LB-100 to reduce the leukemia burden in two mouse models of MLL-AF4+ BCP-ALL: NSG-SEM and NSG-PDX. Strikingly, each drug was individually able to reduce the leukemia burden in the peripheral blood and bone marrow of NSG-SEM mice. Furthermore, acetazolamide as a single agent prolonged the survival of NSG-PDX mice transplanted with leukemic cells derived from an infant patient diagnosed with MLL-AF4+ BCP-ALL. Given that acetazolamide displayed no/mild toxicity towards normal hematopoietic cells and there were no side-effects on mice, this drug could become a valuable addition to the therapeutic regimen for infants with MLL-AF4+ BCP-ALL. Future studies will focus on assessing the effect of each drug over a prolonged period of time (> 2 week treatment), but also in combination with chemotherapeutic agents that are currently used to treat infants with MLL-AF4+ BCP-ALL (e.g. cytarabine, dexamethasone). Given that all three drugs were detrimental to the survival of MLL-AF4+ leukemic cells and disease maintenance *in vivo*, they could become invaluable assets in the clinic to improve the treatment for a class of patients that has seen little improvement over the past 20 years.

## Acknowledgements

This study was supported by a Bloodwise Bennett Senior Fellowship (K.O.) and grants from the Kay Kendall Leukaemia Fund and Cancer Research UK (K.O.). The authors are extremely grateful to patients, their parents and the Bloodwise Childhood Leukaemia Cell Bank for providing MLL-AF4+ samples. The authors would also like to thank Fiona Rossi and Claire Cryer, Bindy Heer and Andrea Corsinotti from the CRM FACS Facility for cell sorting services and flow cytometry advice. We acknowledge the support of James Todd, Andrew Dyer, Allan Booth, Jacek Mendrychowski and Jaimie Kelly from the CRM Animal Facility in animal experimentation. Core facilities at the Edinburgh MRC Centre for Regenerative Medicine were supported by centre grant MR/K017047/1.

## Authorship contribution

C.M. contributed to the design of the study, performed and designed experiments, analyzed the results and wrote the manuscript. A.D., G.C., L.N. and H.J. performed experiments. R.A.A. arranged clinical provision of human fetal tissues. R.S.K provided the PER-494 leukemia cell line. N.A.B and O.P.S. provided the patient material. K.O. conceived and supervised the study and wrote the manuscript. All authors edited and approved the final version of the manuscript for submission.

## Method

### MicroRNA profiling of patients

Primary hematological malignancy samples used in this study were provided by the Blood Cancer UK Childhood Leukaemia Cell Bank. The data were taken from our previous study (Malouf et al. 2021, PMID: 34111240).

### Sorting of human FL hematopoietic cells

Anonymized human fetal livers from morphologically normal 10-20 week-old fetuses were collected following elective medical termination of pregnancy at the Royal Infirmary of Edinburgh after informed written consent (approved by the Lothian Research Ethics Committee, Reference: 08/S1101/1). Fetal livers were dissociated in Flow Cytometry Staining Buffer (ThermoFisher Cat# 00-4222-26) using a 21Gx15mm needle attached to a syringe (BD Microlance Cat# 10472204-X and 3000185) and further CD34-enriched using the CD34 MicroBead Kit Ultrapure human (Millitenyi Cat# 130-100-453) according to manufacturer’s instructions. The CD34+ fraction was stained with the following antibodies (lineage cocktail on APC) in Flow Cytometry Staining Buffer (ThermoFisher Cat# 00-4222-26): APC anti-human CD2 antibody (clone RPA-2.10, Biolegend Cat# 300213), APC anti-human CD3 antibody (clone HIT3, Biolegend Cat# 300311), APC anti-human CD14 antibody (clone M5E2, Biolegend Cat# 301807), APC anti-human CD16 antibody (clone 3G8, Biolegend Cat# 302011), APC anti-human CD56 antibody (clone HCD56, Biolegend Cat# 318309), APC anti-human CD235ab antibody (clone HIR2, Biolegend Cat# 306607), Alexa Fluor^®^ 700 anti-human CD34 antibody (clone 561, Biolegend Cat# 343621), PE anti-human CD45RA antibody (clone HI100, Biolegend Cat# 304107), PE/Cy7 anti-human CD10 antibody (clone eBIOCB-CALLA, ThermoFisher Cat# 25-0106-42), BrilliantViolet™ 421 anti-human CD90 (clone 5E10, Biolegend Cat# 328121), BrilliantViolet™ 510 anti-human CD38 (clone HB7, Biolegend Cat# 356633), BrilliantViolet™ 605 anti-human CD19 (clone HIB19, Biolegend Cat# 302243). The CD34- fraction was stained with the following antibodies in Flow Cytometry Staining Buffer (ThermoFisher Cat# 00-4222-26): Alexa Fluor^®^ 700 antihuman CD34 antibody (clone 561, Biolegend Cat# 343621), APC anti-human CD3 antibody (clone HIT3, Biolegend Cat# 300311), FITC anti-human CD33 antibody (clone P67.6, Biolegend Cat# 366619), PE anti-human CD56 antibody (clone HCD56, Biolegend Cat# 318305), PE/Cy7 anti-human CD20 antibody (clone 2H7, Biolegend Cat# 302311). Cells were incubated on ice for 20 minutes, washed twice with Flow Cytometry Staining Buffer and resuspended in diluted SYTOX AADvanced (ThermoFisher Cat# S10274) to exclude dead cells. Cells were sorted on a BD FACSAria™ II (BD Biosciences) into QIAzol lysis reagent (QIAGEN, Cat# 79306) for RNA extraction. We used the following gating strategy for human hematopoietic stem and progenitor cells: HSC (Lin- CD34+ CD38- CD19- CD45RA- CD90+), MPP (Lin- CD34+ CD38- CD19- CD45RA- CD90-), LMPP (Lin- CD34+ CD38- CD19-CD45RA+), pre-pro-B (Lin- CD34+ CD38+/low CD10- CD19+), pro-B (Lin- CD34+ CD38+/low CD10+ CD19+). We used the following gating strategy for committed and/or mature hematopoietic cells: B-lymphoid cells (CD34- CD20+), NK cells (CD34- CD56+), T-lymphoid cells (CD34- CD3+) and myeloid cells (CD34- CD33+). Reverse transcription and RT-qPCR for microRNA was performed using the TaqMan MicroRNA Reverse Transcription Kit and TaqMan Universal Master Mix II, no UNG according to the manufacturer’s instructions (ThermoFisher Cat# 4366596 and #4440047). MicroRNA expression was assessed using individual Taqman assays (ThermoFisher Assay ID 001973, 000436, 000456, 002198, 002216 and 001973).

### Mice

All animal work was carried out under the regulation of the UK Home Office. Males and females were mated to obtain E14 embryos, with the day of plug detection being counted as day 0 of embryonic development.

### Cell lines

PER-494 (CVCL_A8JU), RS4;11 (ATCC^®^CRL-1873™), SEM (DSMZ ACC 546), Nalm6 (ATCC^®^CRL-3273™), MV4;11 (ATCC^®^CRL-9591 ™), MOLM-13 (DSMZ ACC 554), THP-1 (ATCC**^®^**TIB-200™), Nomo-1 (DSMZ ACC 542) and Kasumi1 (ATCC^®^CRL-2724™) cells were maintained in 20% FCS 1% P/S 1% L-glut RPMI 1640 (cells kindly provided by Professor Mark Dawson, Professor Brian Huntly and Dr. Rishi Sury Kotecha).

### Sorting of E14 fetal liver ckit+ CD34+ and CD45+ CD34- cells

Dissociated fetal liver cells were stained using the following antibody mix in Flow Cytometry Staining Buffer (ThermoFisher Cat# 00-4222-26): APC anti-mouse CD117 (ckit) antibody (clone 2B8, Biolegend Cat# 105811), FITC rat anti-mouse CD34 (clone RAM34, BD Cat# 560238) and PE CD45 monoclonal antibody eBioscience™ (clone 30-F11, ThemoFisher Cat# 12-0451-82). Cells were incubated with the antibodies for 20 minutes on ice, washed twice with Flow Cytometry Staining Buffer and resuspended in diluted SYTOX AADvanced (ThermoFisher Cat# S10274) to exclude dead cells. Cells were sorted on a BD FACSAria™ II (BD Biosciences) into QIAzol lysis reagent (QIAGEN, Cat# 79306) for RNA extraction.

### Sorting of E14 fetal liver E-SLAM HSC

Lineage depletion was performed on dissociated fetal liver cells using the EasySep™ Mouse Hematopoietic Progenitor Cell Isolation Kit (STEMCELL Technologies Cat# 19856) according to the manufacturer’s instructions. Cells were stained using the following antibody mix in Flow Cytometry Staining Buffer (ThermoFisher Cat# 00-4222-26): FITC anti-mouse CD45 antibody (clone 30-F11, Biolegend Cat# 103017), APC anti-mouse CD48 antibody (clone HM48-1, Biolegend Cat# 103411), Pacific Blue™ anti-mouse CD150 antibody (clone TC15-12F12.2, Biolegend Cat# 115923) and PE anti-mouse EPCR monoclonal antibody (clone eBio1560 (1560), ThermoFisher Cat# 12-2012-82). Cells were incubated with the antibodies for 20 minutes on ice, washed twice with Flow Cytometry Staining Buffer and resuspended in diluted SYTOX AADvanced (ThermoFisher Cat# S10274) to exclude dead cells. Cells were sorted on a BD FACSAria™ II (BD Biosciences) into QIAzol lysis reagent (QIAGEN, Cat# 79306) for RNA extraction.

### Electroporation of SEM cells with mimics and siRNA

Transfection of SEM cells with mimics for miR-194, miR-99 and miR-125 the control mimic was performed in RPMI 1640 20% FCS 1% P/S 1% L-glut. We used 200 μL of SEM cells at a concentration of 6 x 10^6^ cells/mL and 300 pmole of *mir*Vana^®^ miRNA mimic (Life technologies Cat#MC10004, MC11021, MC12561) or control (Life technologies Cat#4464058). We used the Gene Pulser Xcell Electroporation system (BioRad Cat# 1652660) with GenePulser/MicroPulser Electroporation Cuvette, 0.4 cm gap (BioRad Cat# 1652088). The following electroporation program was used: Square wave length, 10mseconds, 350 V. After electroporation, cells rested for 15 minutes at room temperature before being diluted 1:20 in RPMI 1640 20% FCS 1% P/S 1% L-glut.

### Lentivirus production in 293T cells and transduction experiments

293T cells (ATCC^®^CRL-3216™) were maintained in 10% FCS, 1% P/S, 1% L-glut DMEM. Lentiviral vectors for the microRNA overexpression were purchased from System Biosciences: Human pre-microRNA Expression Construct Lenti-miR-194-1 (Cat#PMIRH1941PA-1), Human pre-microRNA Expression Construct Lenti-miR-99b (Cat# PMIRH99b-1), Human pre-microRNA Expression Construct Lenti-miR-125a (Cat# PMIRH1125a-PA-1) and empty pMIRH vector (Cat# PMIRH CD511B-1). For the CRISPR-Cas9, we used the following vectors to produce two lentiviruses: pKLV2-U6gRNA5(BbsI)-sEF1aBFP-W (AddGene Cat#67974) and LentiCas9-EGFP (AddGene Cat#63592). We used pMD2.G and psPAX2 as packaging vectors (Addgene Cat# 12259 and 12260). The lentiviral vector, pMD2G and psPAX2 vectors were transfected into HEK293T cells using 1 mg/mL of polyethyleneimine, high molecular weight (Sigma-Aldrich Cat#408727) and serum-free DMEM. The DNA solution was briefly vortexed and allowed to stand at room temperature for 15 minutes before adding to a plate of 60-70% confluent 293T cells. The morning after, transfection media was removed and replaced with fresh media for virus collection at 36-48 hours after transfection. Supernatant containing lentiviral particles was filtered before transduction of fetal liver LSK or leukemia cell lines through a Millex-hV 0.45μm PVDF 33mm Gamma Sterilized filter (Millipore Europe Cat#SLHVM33RS). For leukemia cell lines, supernatant was added directly to cells along with polybrene at a final concentration of 4 μg/mL of polybrene (Santa Cruz Biotechnology Cat# sc-134220). For Mll-AF4+ pMIRH-128a pro-B leukemic blasts, supernatant (MOI > 10) was added on a non-treated plate coated with RetroNectin^®^ Recombinant Human Fibronectin Fragment according to the manufacturer’s instructions (Takara Bio Inc Cat# T100A). Cells were maintained in StemPro^TM^-34 SFM (1X) (ThermoFisher Cat# 10639011) supplemented with 100 ng/mL SCF, 100 ng/mL TPO and 50 ng/mL Flt3 (PeproTech EC Ltd, Cat# 315-14-10, 250-03-10, 250-31L-10) over a 24-hour period. After 3days in culture, cells were collected and used for further studies.

### Cell proliferation, apoptosis assay and *in vitro* drug studies

Cell proliferation assays were conducted in a 6-well plate. Cells were counted Trypan Blue Solution (Life Technologies, Cat# 15250061). Apoptotic cells were detected by double-staining using PE-AnnexinV (Biolegend Cat#640907) and SYTOX Blue Dead Cell Stain (ThermoFisher Cat# S34857) for transduced cells with pMIRH. Apoptotic cells were detected by double-staining using PE-AnnexinV (Biolegend Cat#640907) and SYTOX Red Dead Cell Stain (ThermoFisher Cat# S34859) for transduced cells with pKLV2-U6gRNA5(BbsI)-sEF1aBFP-W (AddGene Cat#67974) and LentiCas9-EGFP Staining was carried out in the Annexin V Binding Buffer according to the manufacturer’s instructions (BD Biosciences Cat# 556454). For drugs studies, we used the following drugs reconstituted in DMSO (Acetazolamide and TACROLIMUS) or sterile water (LB-100): Acetazolamide (Generon Cat# HY-B0782), TACROLIMUS (Cambridge Bioscience Cat# F001) or LB-100 (1.5 mg/kg/day).

### Transplantation of SEM leukemic cells into NSG mice for the rescue experiment and flow cytometry analysis of recipients

SEM leukemic cells that express pMIRH, pMIRH-194, pMIRH-99b or pMIRH-125a-5p were sorted based on GFP expression and tail-vein injected into non-irradiated NSG mice (2500 cells/mouse). Mice were sacrificed once leukemia was established, and single-cell suspensions from peripheral blood and tissues (bone marrow, spleen and liver) were analyzed by flow cytometry using the following antibodies: APC antihuman CD45 (clone 2D1, Biolegend Cat# 368511), PE anti-human CD19 (clone 4G7 Biolegend Cat# 392505), Brilliant Violet 605™ anti-human CD10 (clone HI10a, Biolegend Cat# 312221), Brilliant Violet 421™ anti-human CD33 (clone P67.6, Biolegend Cat# 366621), Brilliant Violet 711™ anti-human IgM (clone MHM-88, Biolegend Cat# 314539) and PE/Cy7 anti-mouse CD45 (clone 30-F11, Biolegend Cat# 103113). Cells were incubated with the antibodies for 20 minutes on ice, washed twice with Flow Cytometry Staining Buffer and resuspended in diluted SYTOX AADvanced (ThermoFisher Cat# S10274) to exclude dead cells. Data were acquired on a BD LSRFortessa™ (BD Biosciences).

### Transplantation of mouse GFP+ Mll-AF4+ pMIRH-128a leukemic cells for the rescue experiment and flow cytometry analysis of recipients

Transplantation recipients (CD45.1/2 mice aged 8-12 weeks old) were irradiated with a total dose of 9.2 Gy (2 doses of 4.6 Gy, 3 hours apart, with a split adaptor). Transduced GFP+ Mll-AF4+ pMIRH-128a cells were were transplanted through tailvein injection (10000 cells/mouse) into irradiated CD45.1/2 recipients along with 20 000 helper bone marrow cells (CD45.1/1). Mice were administered antibiotics after transplantation through their drinking water (0.1 mg/mL enrofloxacin, 10% Batyril solution from Bayer) and bled on a monthly basis. Blood counts were measured on a Celltac MEK-6500K (Nihon Kohden). For flow cytometry analysis of peripheral blood and tissues, red blood cell lysis was carried out with BD Pharm Lyse™ lysing solution according to the manufacturer’s instructions (BD Biosciences Cat# 555899). Cells were stained in Flow Cytometry Staining Buffer using the following mixture of antibodies: APC-eFluor 780-CD45.2 monoclonal antibody eBioscience™ (clone 104, ThermoFisher Cat# 47-0454-82), PE-CD45.1 monoclonal antibody eBioscience™ (clone A20, ThermoFisher Cat# 12-0543-83), eFluor450-CD11b monoclonal antibody eBioscience™ (clone M1/70, ThermoFisher Cat# 48-0112-80), Alexa Fluor^®^ 700 anti-mouse Ly-6G/Ly-6C (Gr-1) antibody (clone RB6-8C5, Biolegend Cat# 108422), PE/Cy7 anti-mouse/human CD45R/B220 antibody (clone RA3-6B2, Biolegend Cat# 103222), Brilliant Violet 605™ anti-mouse CD19 antibody (clone 6D5, Biolegend Cat# 115539), APC-IgM mouse monoclonal antibody (clone II/41, ThermoFisher Cat# 17-5790-82), Brilliant Violet 711™-CD3 anti-mouse antibody (clone 17A2, Biolegend Cat# 100241). We also used the following antibodies to detect B-cells and LSK cells: Alexa Fluor^®^ 700 anti-mouse/human CD45R/B220 (clone RA3-6B2, Biolegend, Cat# 103231), Brilliant Violet 421™ antimouse CD117 (c-kit) antibody (clone 2B8, Biolegend, Cat# 105827), CD127 (IL7R) monoclonal antibody, PE, eBioscience™ (clone A7T34, ThermoFisher Cat# 12-1271-82), APC anti-mouse CD3ε antibody (clone I45-2C11, Biolegend Cat# 100312), APC anti-mouse TER-119 antibody (clone TER119, Biolegend Cat# 116212), APC anti-mouse F4/80 antibody (clone BM8, Biolegend Cat# 123116), APC anti-mouse Nk1.1 antibody (clone PK136, Biolegend Cat# 108709), APC antimouse Ly-6G/Ly-6C (Gr-1) antibody (clone RB6-8C5, Biolegend Cat# 108412), APC anti-mouse/human CD45R/B220 antibody (clone RA3-6B2, Biolegend Cat# 103211), APC anti-mouse CD19 antibody (clone 6D5, Biolegend, Cat#115511), Brilliant Violet 421™ anti-mouse CD117 (ckit) antibody (clone 2B8, Biolegend, Cat# 105827), PE/Cy7 anti-mouse Ly-6A/E (Sca1) antibody (clone E13-161.7, Biolegend Cat# 122513), CD127 (IL7R) monoclonal antibody, PE, eBioscience™ (clone A7T34, ThermoFisher Cat# 12-1271-82), CD135 (Flt3) monoclonal antibody, biotin, eBioscience™ (clone A2F10, ThermoFisher Cat# 13-1351-81). Cells were incubated on ice for 20 minutes, washed twice with Flow Cytometry Staining Buffer and resuspended in diluted SYTOXAADvanced (ThermoFisher, Cat# S10274) to exclude dead cells. Data were acquired on a BD LSRFortessa™ (BD Biosciences).

### Drug studies in NSG mice using SEM cells and primary patient sample

100 000 leukemic cells were transplanted by tail-vein injection into non-irradiated NSG mice. 3-4 weeks post-transplant, once we detected human cells in the peripheral blood, we started the 12-day treatment protocol. Day 1 is considered the first day of the treatment (first drug injection). On odd days, mice received intraperitoneal injections of Acetazolamide (2 x 20 mg/kg/day), TACROLIMUS (1 x 10 mg/kg/day) or LB-100 (1 x 1.5 mg/kg/day). On even days, mice were bled to monitor disease progression (NSG-SEM only). On day 12, all NSG-SEM mice were sacrificed for extensive post-mortem analysis. For NSG-PDX mice, vehicle and Acetazolamide treated mice were sacrificed when a full-blown leukemia had developed.

### Quantitative PCR for mRNA

RNA extraction and reverse transcription were done using the RNeasy Micro Kit (QIAGEN Cat# 74004). For reverse transcription of mRNA, we used the iScript Ready-to-Use cDNA Supermix of iScript Advanced cDNA synthesis kit for RT-qPCR (Bio-Rad Laboratories Ltd Cat# 1708841 or Cat#1725037) according to the manufacturer’s instructions. Primers were designed using Primer3 and tested. Primer sequences can be found in Supplementary Table S1. We used the Brilliant III Ultra-Fast SYBR^®^ Green qPCR Master Mix according to the manufacturer’s instructions (Agilent Cat# 600883). Data were acquired on a QuantStudio™ 7 Flex Real-Time PCR System (ThermoFisher).

### Western blot

Protein extraction was carried out in RIPA buffer (50 mM sodium chloride, 1% NP-40, 0.5% sodium deoxycholate, 0.1% SDS, 50 mM Tri pH 8.0) supplemented with a protease inhibitor cocktail tablet (Roche Cat# 04693116001). Cell pellets were washed twice in PBS, resuspended in complete RIPA buffer and incubated on ice for 30 minutes. Genomic DNA was fragmented using a needle/syringe. Protein quantification was performed using the Lowry assay according to manufacturer’s instruction (BioRad Cat# 5000112). 5-10 μg of total protein extract was denaturated (95°C for 5 minutes) into 1X Laemmli buffer (Sigma Cat# S3401-10VL) and loaded on a homemade gel (12% separation: 12% Acrylamide (37.5:1), 625 mM Tris pH 8.8, 0.17% SDS, 0.08% TEMED, 0.8% APS. 4% stacking: 4% Acrylamide (37.5:1), 120 mM Tris pH 6.8, 0.1% SDS, 0.01% TEMED, 0.4% APS). Migration was carried out in Tris-Glycine/SDS running buffer (25 mM Tris base, 190 mM glycine, 0.1% SDS), followed by wet protein transfer to a PVDF membrane in transfer buffer (25 mM Tris base, 190 mM glycine, 10% methanol). Membranes were blocked in 5% milk TBST Buffer for one hour at 4°C, with mild shaking. Primary antibody was added at a dilution of 1:1000 in 5% milk TBST Buffer and incubated overnight at 4°C, with mild shaking: human Carbonic Anhydrase VB/CA5B antibody (Biotechne Cat#AF3176), rabbitt anti-PPP2R5C antibody affinity purified (Bethyl Laboratories Inc Cat#A308-813A-T), PPP3CA polyclonal antibody(Elabscience Cat# E-AB-14813), monoclonal anti-β-actin antibody produced in mouse(Sigma Cat# A2228). Membranes were washed 3 times for 5 minutes in TBTS, followed by incubation with a HRP-conjugated secondary antibody (ThermoFisher, Cat# A16066 and A16096). Membranes were washed 3 times for 5 minutes in TBST and 1 time for 5 minutes in TBS. Protein signal was detected using the Clarity™ Western ECL Substrate (BioRad Cat# 1705061) and image acquisition on the ChemiDoc Imaging System (BioRad Cat# 17001401). Image analysis and quantification was performed on Image Lab Software for PC Version 6.1 (BioRad, SOFT-LIT-170-9690-ILSPC-V-6-1). Relative protein quantification for each protein was done according to a reference band (control siRNA), and the signal for the protein of interest (CA5B, PPP3CA or PPP2R5C) was adjusted with the relative quantification of the beta-actin signal. The beta-actin signal was acquired from the same membrane as the protein of interest. Membranes were rinsed briefly in TBS after each experiment and dried overnight at room temperature. The second staining of blot membranes was done 2-3 weeks later, once the signal from the previous antibody had disappeared.

### Cloning of 3’UTR into pGL3promoter vector and guide RNA into the pKLV2-U6gRNA5(BbsI)-sEF1aBFP-W vector

The *CA5B, PPP3CA and PPP2R5C* UTR were amplified by PCR and cloned in the XbaI site of the pGL3-promoter vector (Promega Cat# E1741). Guide RNAs were designed using CHOPCHOP (https://chopchop.cbu.uib.no/) and cloned into the Bbs1 site of the pKLV2-U6gRNA5(BbsI)-sEF1aBFP-W vector. Oligo annealing for guide RNAs was done using the following method: 37 °C, 30min → 95 °C, 50min → ramp down to 25 °C at 5 °C/min. XbaI or Bbs1 digestion (New England Biolabs Cat# R0145S or R0539S), DNA gel extraction (QIAGEN Cat# 28704) and ligation (New England Biolabs Cat# M0202S) were conducted according to the manufacturer’s instructions. Bacterial transformation was carried out in One Shot™ Stbl3™ Chemically Competent E.coli according to the manufacturer’s instruction (ThermoFisher Cat# C737303). Plasmids were purified with the QIAGEN Plasmid MiniPrep, Midiprep or Maxiprep kits (QIAGEN Cat# 27104, 12643 and 12662).

### Luciferase assay

On the day of the transfection, HEK293T cells were harvested and seeded in DMEM 10% FCS 1%P/S 1%L-glut at a concentration of 100 000 cells/well in a 96-well solid white plate (Fisher Scientific Cat# 10022561). The transfection mix for each well consisted of 98 ng pGL3-promoter, 100 ng pMIRH and 2 ng pRL-SV40 (Promega Cat# E2231) in 100 uL of serum-free DMEM and 1 mg/mL of polyethyleneimine, high molecular weight (Sigma-Aldrich Cat#408727). The DNA solution was briefly vortexed and allowed to stand at room temperature for 15 minutes before adding to a plate. 48 hours after transfection, the Firefly and Renilla luciferase activity were measured using the Dual-Glo Luciferase Assay System (Promega Cat# E2920) and a GloMax Discover MicroPlate Reader (Promega Cat# GM3000) according to the manufacturer’s instructions.

### MicroRNA target prediction and data analysis

TargetScan ^38^ and PicTar ^39^ were used to generate a list of *in silico* target genes for miR-130b and miR-128a in human and mouse. The cross-reference of target genes between tool and species was done using Venny (http://bioinfogp.cnb.csic.es/tools/venny/). Graphs were generated using GraphPad Prism 6 (GraphPad software corporation) with statistical tests specified in the figure legends.

