## Supplemental Material for "A microRNA expression signature in infant t(4;11) MLL-AF4+ BCP-ALL uncovers novel therapeutic targets"

- Supplementary Figure Legends
- Supplementary Table S1

#### Supplemental Figure Legends

**Supplemental Figure 1. MiR-194, miR-99b and miR-125a-5p are all downregulated in t(4;11) MLL-AF4 BCP-ALL patients.** (A) MiR-194, (B) miR-99b and (C) miR-125a-5p expression in normal mouse FL and adult bone marrow hematopoietic stem cells (Lineage- CD150+ CD48- EPCR+), hematopoietic progenitors (ckit+CD34+) and differentiated hematopoietic cells (CD45+CD34-). MIR-128a, miR-130b, miR-194, miR-99b and miR-125a-5p expression in (D) SEM and (E) RS4;11 leukemia cell lines. Data are presented as Mean ± SEM and compared using a Mann-Whitney U test with bilateral p-value: $p < 0.05$ (*), $p < 0.01$ (**), $p < 0.001$ (***) and $p < 0.0001$ (****).

**Supplemental Figure 2. MiR-194, miR-99b and miR-125a-5p impair MLL-AF4+ leukemia maintenance.** GFP chimerism in the (A) spleen, (B) peripheral blood, (D) liver and (D) lungs of Mll-AF4+ pMIRH-128a BCP-ALL mice that overexpress miR-194, miR-99b or miR-125a-5p. (E) Proportion of CKIT$^{high}$IL7R+ and CKIT$^{low}$IL7R+ leukemic blasts in the spleen of Mll-AF4+ pMIRH-128a BCP-ALL mice that overexpress miR-194, miR-99b or miR-125a-5p. (F) RT-qPCR to confirm the overexpression of miR-194, miR-99b or miR-125a-5p in the bone marrow of rescue mice (end of experiment/leukemia). Data are presented as Mean ± SEM and compared using a Mann-Whitney U test with bilateral p-value: $p < 0.05$ (*), $p < 0.01$ (**), $p < 0.001$ (***) and $p < 0.0001$ (****).

**Supplemental Figure 3. MiR-194, miR-99b and miR-125a-5p act through their downstream targets CA5B, PPP3CA and PPP2R5C.** (A) *CA5B*, (B) *PPP3CA* and (C) *PPP2R5C* expression in MLL-AF4+ BCP-ALL infant patient and cord blood B-cell progenitors (CB BCP) (GSE79450). (D) *CA5B*, (E) *PPP3CA* and (F) *PPP2R5C* expression in leukemia cell lines. (G) *Ca5b*, (H) *Ppp3ca* and (I) *Ppp2r5c* expression in Mll-AF4+ pMIRH-128a pre-transplant cells (GFP+ pro-B cells transduced with pMIRH, pMIRH-194, pMIRH-99b or pMIRH-125a-5p). (J) Western blots of SEM cells transfected with miRVANA® mimics for CA5B, PPP3CA and PPP2R5C. (K) *CA5B*, (L) *PPP3CA* and (M) *PPP2R5C* expression in SEM cells transduced with Cas9-GFP and KPL474-guideRNA-BFP. RT-qPCR (top) and western blot (low) are shown. Data are presented as Mean ± SEM and compared using a Mann-Whitney U test with bilateral p-value: $p < 0.05$ (*), $p < 0.01$ (**), $p < 0.001$ (***) and $p < 0.0001$ (****).

**Supplemental Figure 4. Acetazolamide, Tacrolimus and LB-100 impair the survival of MLL-AF4+ pro-B leukemic blasts while having minimal effect on normal FL and BM mouse LSK.** Proliferation of (A) PER494, (B) RS4;11 and (C) MV4;11 leukemia cells exposed to Acetazolamide, Tacrolimus and LB-100 (10 µM). (D) Fold change in AnnexinV+ cells of human FL CD34+ cells exposed to Acetazolamide, Tacrolimus and LB-100 (10 µM) (n = 2). Data are presented as Mean ± SEM and compared using a Mann-Whitney U test with bilateral p-value: $p < 0.05$ (*), $p < 0.01$ (**), $p < 0.001$ (***) and $p < 0.0001$ (****).

**Supplemental Figure 5. Acetazolamide, Tacrolimus and LB-100 impair the maintenance of MLL-AF4+ pro-B ALL.** Weight fluctuation of NSG-SEM mice over the drug treatment: (A) Vehicle, (B) Acetazolamide, (C) Tacrolimus and (D) LB-100. (E) Spleen and (F) liver sizes of NSG-SEM vehicle and drug treated mice. (G) Spleen and (H) liver sizes of NSG-PDX vehicle and Acetazolamide mice. Data are presented as Mean ± SEM and compared using a Mann-Whitney U test with bilateral p-value: $p < 0.05$ (*), $p < 0.01$ (**), $p < 0.001$ (***) and $p < 0.0001$ (****).

#### Supplemental Table S1. Primers

| Name | Species | Sequence (5' to 3') |
|---|---|---|
| MLL-AF4.F (RT-qPCR) | Human | ACAGAAAAAAGTGGCTCCCCG |
| MLL-AF4.R (RT-qPCR) | Human | TATTGCTGTCAAAGGAGGCGG |
| CA5B.F (RT-qPCR) | Human | CTTCAAGCCTCTCCAGGCAA |
| CA5B.R (RT-qPCR) | Human | CCAGAGTGGATGCAAGGCTC |
| PPP3CA.F (RT-qPCR) | Human | GGAGATGTCCGAGCCCAAG |
| PPP3CA.R (RT-qPCR) | Human | TGGAACAGCTTTCACCACCC |
| PPP2R5C.F (RT-qPCR) | Human | AATAAAGCGGGCAGCAGGAT |
| PPP2R5C.R (RT-qPCR) | Human | TCAGCAGGAGGAACATCTCG |
| hCA5B.Guide1.F (cloning) | Human | CACCGTCCGAGATTCATGCCAGCGAGG |
| hCA5B.Guide1.R (cloning) | Human | AAACCCTCGCTGGCATGAATCTCGGAC |
| hCA5B.Guide2.F (cloning) | Human | CACCGGAACAGTGCGATTCATCAGTGG |
| hCA5B.Guide2.R (cloning) | Human | AAACCCACTGATGAATCGCACTGTTCC |
| hPPP3CA.Guide2.F (cloning) | Human | CACCGTTGTCGACGACCGACAGGGTGG |
| hPPP3CA.Guide2.R (cloning) | Human | AAACCCACCCTGTCGGTCGTCGACAAC |
| hPPP3CA.Guide3.F (cloning) | Human | CACCGCGACCGACAGGGTGGTGAAAGG |
| hPPP3CA.Guide3.R (cloning) | Human | AAACCCTTTCACCACCCTGTCGGTCGC |
| hPPP2R5C.Guide1.F (cloning) | Human | CACCGACGGGAGCGGAATTTGACCCGG |
| hPPP2R5C.Guide1.R (cloning) | Human | AAACCCGGGTCAAATTCCGCTCCCGTC |
| hPPP2R5C.Guide3.F (cloning) | Human | CACCGTCCGCTCCCGTAGGATTGGAGG |
| hPPP2R5C.Guide3.R (cloning) | Human | AAACCCTCCAATCCTACGGGAGCGGAC |
| hCA5B-UTR-XbaI.F | Human | CATGTCTAGAAGGCTGGTCTTGAACTCCTG |
| hCA5B-UTR-XbaI.R | Human | CATGTCTAGAGGGAAGACGGGATAGCTCTT |
| hPPP3CA-UTR-XbaI.F | Human | CATGTCTAGACTGTTCTGTCATGTGCCACA |
| hPPP3CA-UTR-XbaI.R | Human | CATGTCTAGACGGACTCAGTGGAAGGATGA |
| hPPP2R5C-UTR-XbaI.F | Human | CATGTCTAGAGCGAGGATGACGTTTAGTGG |
| hPPP2R5C-UTR-XbaI.R | Human | CATGTCTAGAGTACCAGAGCTGCACAAGGA |

Figure S1

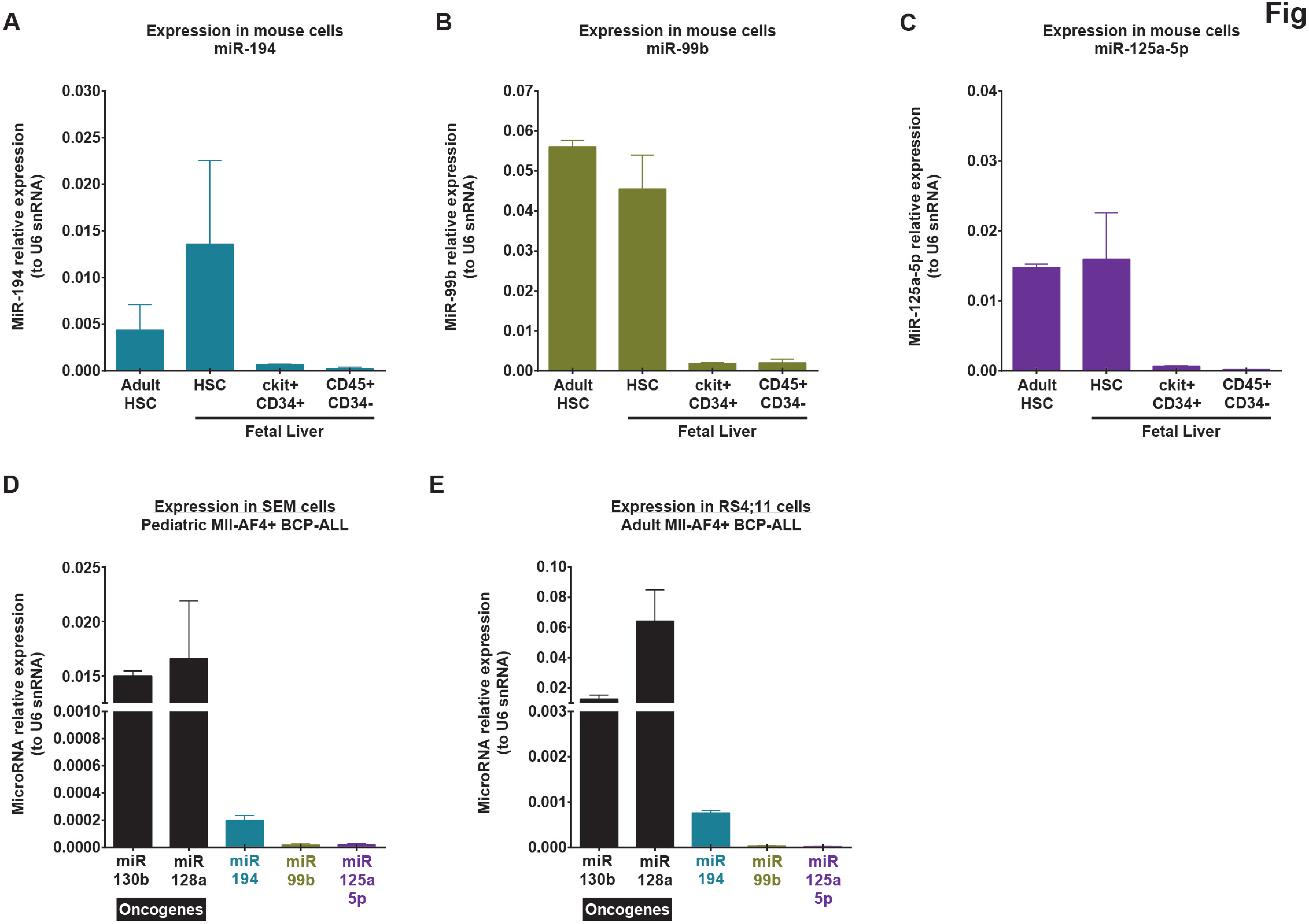
A
Expression in mouse cells
miR-194
MiR-194 relative expression (to U6 snRNA)
0.030
0.025
0.020
0.015
0.010
0.005
0.000
Adult HSC
HSC
ckit+ CD34+
CD45+ CD34-
Fetal Liver
B
Expression in mouse cells
miR-99b
MiR-99b relative expression (to U6 snRNA)
0.07
0.06
0.05
0.04
0.03
0.02
0.01
0.00
Adult HSC
HSC
ckit+ CD34+
CD45+ CD34-
Fetal Liver
C
Expression in mouse cells
miR-125a-5p
MiR-125a-5p relative expression (to U6 snRNA)
0.04
0.03
0.02
0.01
0.00
Adult HSC
HSC
ckit+ CD34+
CD45+ CD34-
Fetal Liver
D
Expression in SEM cells
Pediatric Mll-AF4+ BCP-ALL
MicroRNA relative expression (to U6 snRNA)
0.025
0.020
0.015
0.0010
0.0008
0.0006
0.0004
0.0002
0.0000
miR 130b
miR 128a
miR 194
miR 99b
miR 125a 5p
Oncogenes
E
Expression in RS4;11 cells
Adult Mll-AF4+ BCP-ALL
MicroRNA relative expression (to U6 snRNA)
0.10
0.08
0.06
0.04
0.02
0.003
0.002
0.001
0.000
miR 130b
miR 128a
miR 194
miR 99b
miR 125a 5p
Oncogenes

### Figure S2

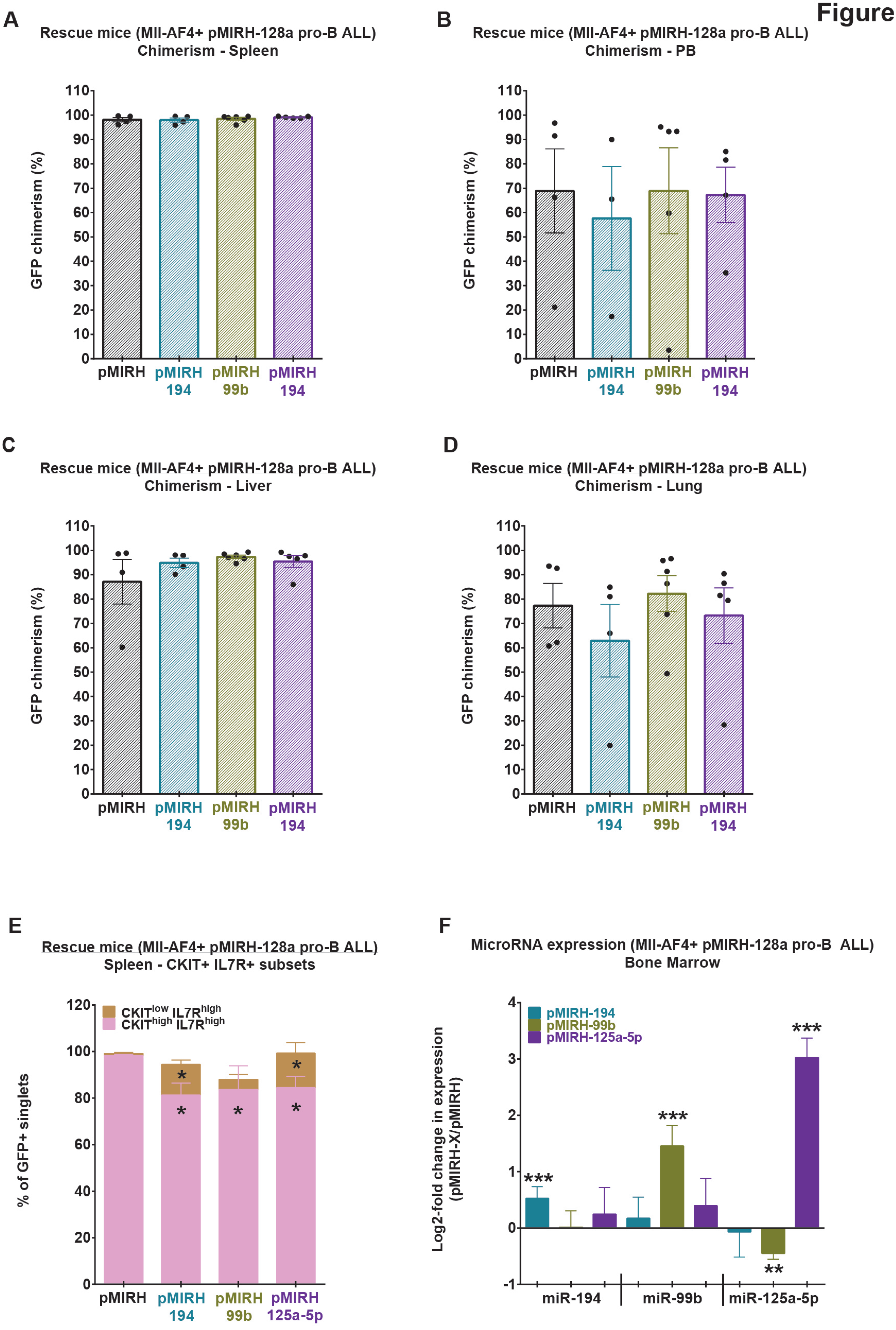

A
Rescue mice (Mll-AF4+ pMIRH-128a pro-B ALL)
Chimerism - Spleen
GFP chimerism (%)
pMIRH
pMIRH 194
pMIRH 99b
pMIRH 194
B
Rescue mice (Mll-AF4+ pMIRH-128a pro-B ALL)
Chimerism - PB
GFP chimerism (%)
pMIRH
pMIRH 194
pMIRH 99b
pMIRH 194
C
Rescue mice (Mll-AF4+ pMIRH-128a pro-B ALL)
Chimerism - Liver
GFP chimerism (%)
pMIRH
pMIRH 194
pMIRH 99b
pMIRH 194
D
Rescue mice (Mll-AF4+ pMIRH-128a pro-B ALL)
Chimerism - Lung
GFP chimerism (%)
pMIRH
pMIRH 194
pMIRH 99b
pMIRH 194
E
Rescue mice (Mll-AF4+ pMIRH-128a pro-B ALL)
Spleen - CKIT+ IL7R+ subsets
CKIT^low IL7R^high
CKIT^high IL7R^high
% of GFP+ singlets
pMIRH
pMIRH 194
pMIRH 99b
pMIRH 125a-5p
F
MicroRNA expression (Mll-AF4+ pMIRH-128a pro-B ALL)
Bone Marrow
pMIRH-194
pMIRH-99b
pMIRH-125a-5p
Log2-fold change in expression (pMIRH-X/pMIRH)
miR-194
miR-99b
miR-125a-5p

Figure S3

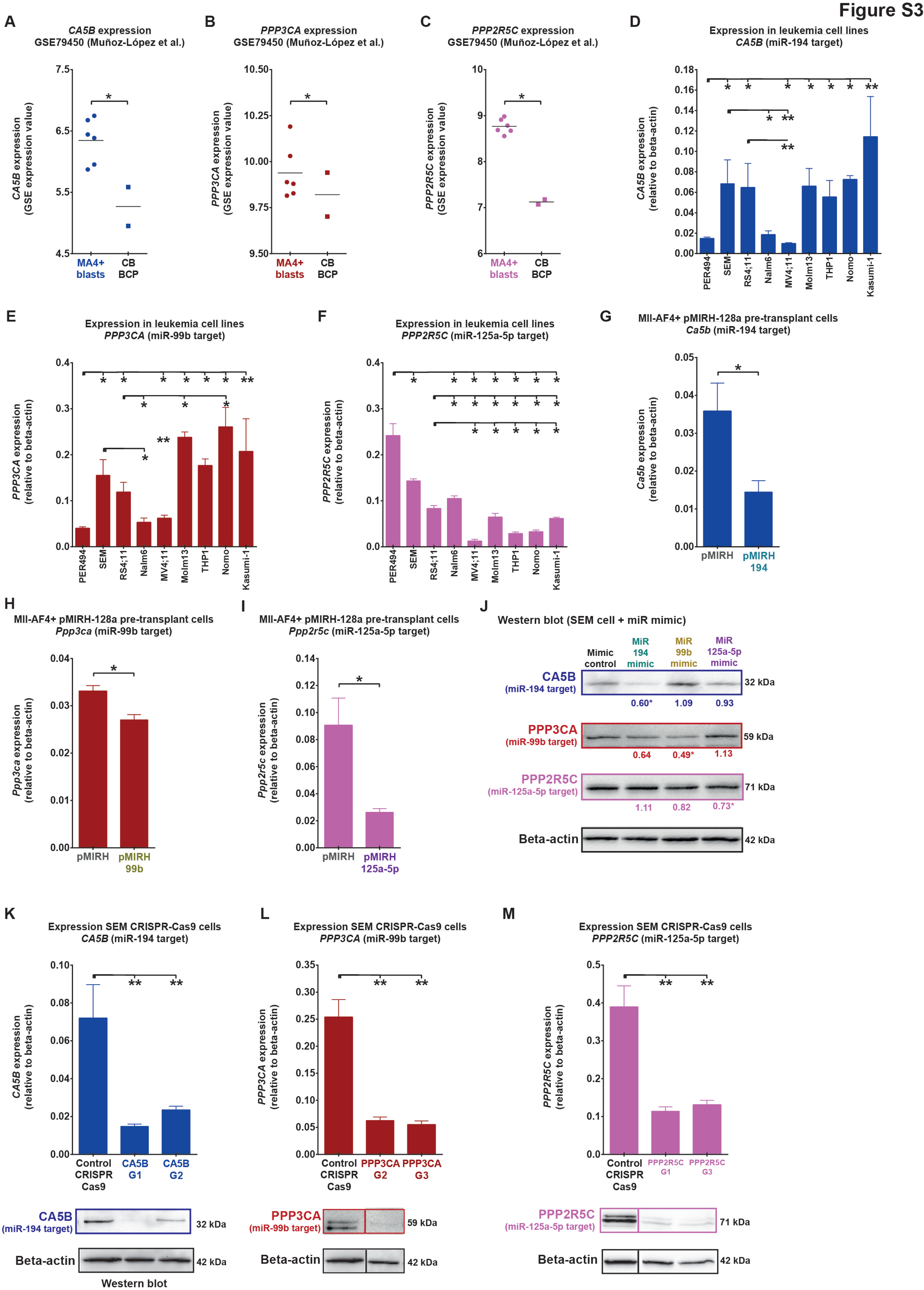
A
CA5B expression
GSE79450 (Muñoz-López et al.)
CA5B expression (GSE expression value)
MA4+ blasts
CB BCP
B
PPP3CA expression
GSE79450 (Muñoz-López et al.)
PPP3CA expression (GSE expression value)
MA4+ blasts
CB BCP
C
PPP2R5C expression
GSE79450 (Muñoz-López et al.)
PPP2R5C expression (GSE expression value)
MA4+ blasts
CB BCP
D
Expression in leukemia cell lines
CA5B (miR-194 target)
CA5B expression (relative to beta-actin)
PER494
SEM
RS4;11
Nalm6
MV4;11
Molm13
THP1
Nomo
Kasumi-1
E
Expression in leukemia cell lines
PPP3CA (miR-99b target)
PPP3CA expression (relative to beta-actin)
F
Expression in leukemia cell lines
PPP2R5C (miR-125a-5p target)
PPP2R5C expression (relative to beta-actin)
G
Mll-AF4+ pMIRH-128a pre-transplant cells
Ca5b (miR-194 target)
Ca5b expression (relative to beta-actin)
pMIRH
pMIRH 194
H
Mll-AF4+ pMIRH-128a pre-transplant cells
Ppp3ca (miR-99b target)
Ppp3ca expression (relative to beta-actin)
pMIRH
pMIRH 99b
I
Mll-AF4+ pMIRH-128a pre-transplant cells
Ppp2r5c (miR-125a-5p target)
Ppp2r5c expression (relative to beta-actin)
pMIRH
pMIRH 125a-5p
J
Western blot (SEM cell + miR mimic)
Mimic control
MiR 194 mimic
MiR 99b mimic
MiR 125a-5p mimic
CA5B (miR-194 target)
0.60* 1.09 0.93
32 kDa
PPP3CA (miR-99b target)
0.64 0.49* 1.13
59 kDa
PPP2R5C (miR-125a-5p target)
1.11 0.82 0.73*
71 kDa
Beta-actin
42 kDa
K
Expression SEM CRISPR-Cas9 cells
CA5B (miR-194 target)
CA5B expression (relative to beta-actin)
Control CRISPR Cas9
CA5B G1
CA5B G2
CA5B (miR-194 target)
32 kDa
Beta-actin
42 kDa
Western blot
L
Expression SEM CRISPR-Cas9 cells
PPP3CA (miR-99b target)
PPP3CA expression (relative to beta-actin)
Control CRISPR Cas9
PPP3CA G2
PPP3CA G3
PPP3CA (miR-99b target)
59 kDa
Beta-actin
42 kDa
M
Expression SEM CRISPR-Cas9 cells
PPP2R5C (miR-125a-5p target)
PPP2R5C expression (relative to beta-actin)
Control CRISPR Cas9
PPP2R5C G1
PPP2R5C G3
PPP2R5C (miR-125a-5p target)
71 kDa
Beta-actin
42 kDa

### Figure S4

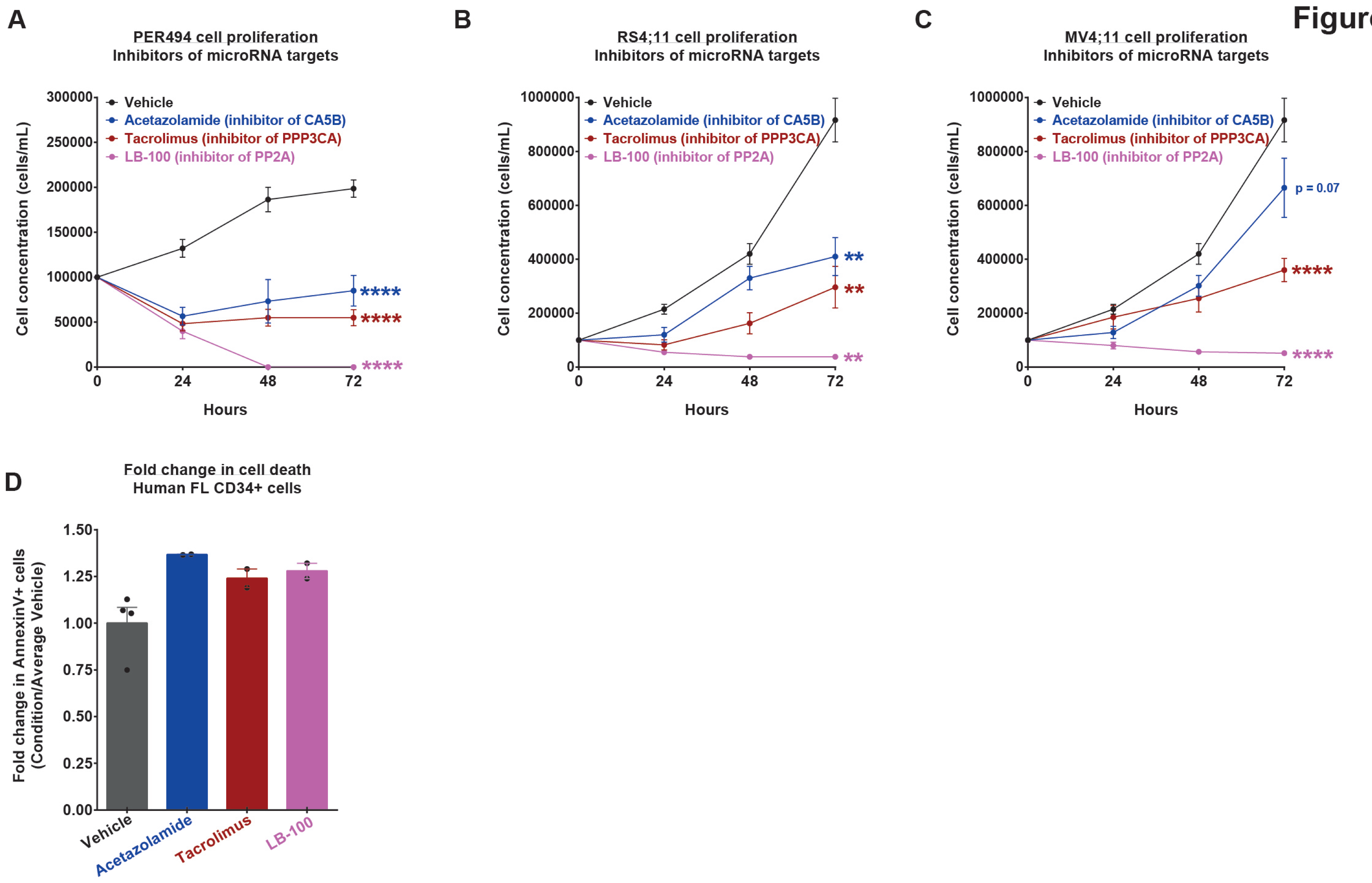
A
PER494 cell proliferation
Inhibitors of microRNA targets
Cell concentration (cells/mL)
300000
250000
200000
150000
100000
50000
0
Vehicle
Acetazolamide (inhibitor of CA5B)
Tacrolimus (inhibitor of PPP3CA)
LB-100 (inhibitor of PP2A)
****
****
****
0
24
48
72
Hours
B
RS4;11 cell proliferation
Inhibitors of microRNA targets
Cell concentration (cells/mL)
1000000
800000
600000
400000
200000
0
Vehicle
Acetazolamide (inhibitor of CA5B)
Tacrolimus (inhibitor of PPP3CA)
LB-100 (inhibitor of PP2A)
**
**
**
0
24
48
72
Hours
C
MV4;11 cell proliferation
Inhibitors of microRNA targets
Cell concentration (cells/mL)
1000000
800000
600000
400000
200000
0
Vehicle
Acetazolamide (inhibitor of CA5B)
Tacrolimus (inhibitor of PPP3CA)
LB-100 (inhibitor of PP2A)
p = 0.07
****
****
0
24
48
72
Hours
D
Fold change in cell death
Human FL CD34+ cells
Fold change in AnnexinV+ cells
(Condition/Average Vehicle)
1.50
1.25
1.00
0.75
0.50
0.25
0.00
Vehicle
Acetazolamide
Tacrolimus
LB-100

### Figure S5

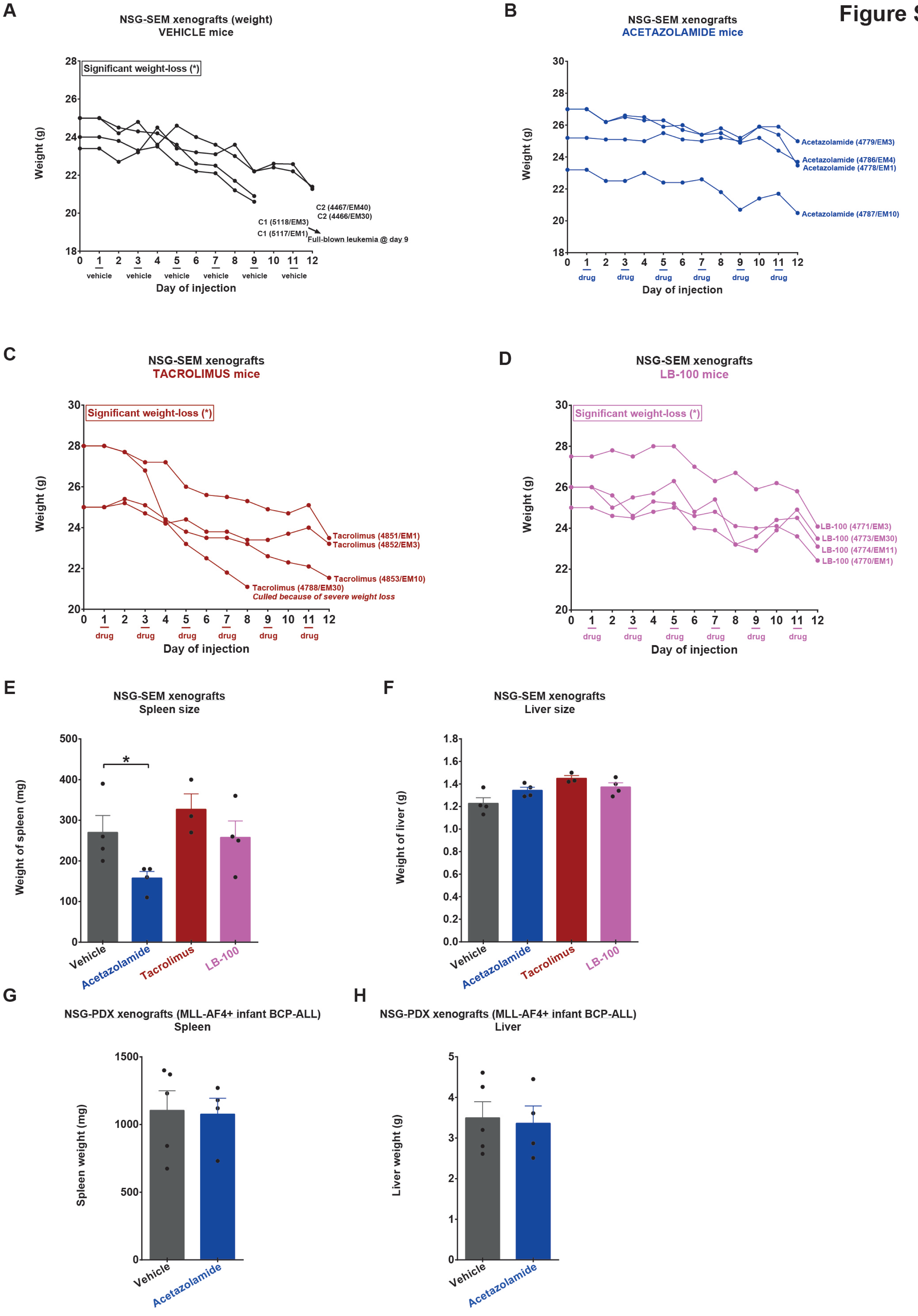
A
NSG-SEM xenografts (weight)
VEHICLE mice
Significant weight-loss (*)
Weight (g)
C2 (4467/EM40)
C2 (4466/EM30)
C1 (5118/EM3)
C1 (5117/EM1)
Full-blown leukemia @ day 9
vehicle
Day of injection
B
NSG-SEM xenografts
ACETAZOLAMIDE mice
Acetazolamide (4779/EM3)
Acetazolamide (4786/EM4)
Acetazolamide (4778/EM1)
Acetazolamide (4787/EM10)
drug
C
NSG-SEM xenografts
TACROLIMUS mice
Tacrolimus (4851/EM1)
Tacrolimus (4852/EM3)
Tacrolimus (4853/EM10)
Tacrolimus (4788/EM30)
Culled because of severe weight loss
D
NSG-SEM xenografts
LB-100 mice
LB-100 (4771/EM3)
LB-100 (4773/EM30)
LB-100 (4774/EM11)
LB-100 (4770/EM1)
E
NSG-SEM xenografts
Spleen size
Weight of spleen (mg)
*
Vehicle
Acetazolamide
Tacrolimus
LB-100
F
NSG-SEM xenografts
Liver size
Weight of liver (g)
G
NSG-PDX xenografts (MLL-AF4+ infant BCP-ALL)
Spleen
Spleen weight (mg)
H
NSG-PDX xenografts (MLL-AF4+ infant BCP-ALL)
Liver
Liver weight (g)
